# Time-resolved tmFRET reveals GTP-coupled conformational changes in Mfn1

**DOI:** 10.1101/2025.10.08.681278

**Authors:** S.M. Hurwitz, W.N. Zagotta, S.E. Gordon, S Hoppins

**Author notes:** Corresponding author: Suzanne Hoppins, University of Washington, Department of Biochemistry, 1959 NE Pacific Street, Health Sciences Building, J383, Seattle, WA 98195.

## Abstract

Outer mitochondrial membrane fusion is mediated by the mitofusin paralogs Mfn1 and Mfn2. Nucleotide-driven self-assembly and conformational changes are required for regulated membrane fusion activity, but the allosteric mechanisms remain enigmatic due to incomplete structural information. In this study, we investigate the GTP-coupled conformational dynamics of Mfn1 using time-resolved transition metal ion fluorescence resonance energy transfer (tmFRET). Using the minimal Mfn1 construct with the GTPase domain and helical bundle 1 (HB1) connected by Hinge 2, we engineered FRET pairs by incorporating a fluorescent noncanonical amino acid donor and a metal ion acceptor. For each state of the catalytic cycle, we measured tmFRET with fluorescence lifetimes and determined distance distributions, which can capture complex structural heterogeneity. Our distance measurements for the GDP-bound state matched predictions from the atomic resolution structure, establishing that the same open state, with GTPase and HB1 domains far apart, exists in solution. Our data reveal that the transition state is not a single closed state with HB1 stably contacting the GTPase domain. Rather, the distance distributions indicate that the presence of GDP + Pi results in an equilibrium between the open and closed states. We also captured the GTP-bound and nucleotide-free states of Mfn1. GTP binding favors the open state and the conformation of the apo state is distinct from either nucleotide bound state. Together, these findings redefine our understanding of GTP-driven conformational dynamics in Mfn1, demonstrating an unexpected conformational reversal in a single catalytic cycle and a heterogeneous transition-state ensemble with implications for the mechanism and regulation of mitochondrial membrane fusion.

**eTOC summary:** Hurwitz et al. leverage a novel FRET-based approach to quantify conformational dynamics of Mfn1, a member of the dynamin superfamily that mediates mitochondrial outer membrane fusion. Their findings shed light on how GTP hydrolysis governs the conformational state of Mfn1, as well as the energetics underlying a key conformational transition.

## Introduction

Eukaryotic cells are functionally compartmentalized by membrane bound organelles. These organelle membranes are dynamic, and ongoing fusion and fission events play essential roles in protein and lipid trafficking as well as in quality control mechanisms. The fusion and fission of diverse cellular membranes are mediated by the dynamin superfamily of proteins (DSPs)(Hutson et al.; Khurana and Pucadyil, 2023). DSPs are mechanochemical enzymes that hydrolyze GTP. DSPs have shared domain organization, including a globular GTPase domain, helical bundle domains, and a lipid-interacting/targeting domain. Universally, GTP allosterically regulates conformational changes in DSPs to drive oligomerization and membrane remodeling (Hutson et al.; Kalia and Frost, 2019; Jimah and Hinshaw, 2019).

Mechanistic understanding of the DSP family is largely derived from studies of dynamin and dynamin-related protein 1 (Drp1). Both utilize mechano-constriction to catalyze fission of their target membranes (plasma membrane and mitochondria, respectively) (Helle et al., 2017; Alimohamadi et al., 2025). Our mechanistic understanding of fission DSPs has benefited from the accumulation of structural information that captures different conformational states, although allosteric regulation at the individual protein level remains largely undetermined (Hutson et al.; Kalia and Frost, 2019; Chappie et al., 2011; Chappie and Dyda, 2013; Cocucci et al., 2014).

Less is known about the mechanism of fusion DSPs, including the mitofusin paralogs, Mfn1 and Mfn2. The mitofusins are localized to the mitochondrial outer membrane and together mediate fusion of this membrane (Hoppins et al., 2011; Chen et al., 2003; Santel et al., 2003). In concert, mitochondrial fusion, fission and microtubule-based transport regulate mitochondrial size, shape, and distribution throughout the cell (Tilokani et al., 2018; Tábara et al., 2025; Mattie et al., 2019; Suomalainen and Nunnari, 2024; Quintana-Cabrera and Scorrano, 2023; Dorn, 2019). These dynamic properties are also essential to maintain and modulate mitochondrial functions such as metabolism, innate immune response, and apoptosis. Dysregulation of mitochondrial fusion resulting from mutations in *MFN2* causes the neurodegenerative disease Charcot Marie Tooth Type 2A (CMT2A) (Züchner et al., 2004; Feely et al., 2011; Alberti et al., 2024). Other neurogenerative diseases including ALS, Alzheimer’s, and Parkinson’s disease are also associated with aberrant mitochondrial dynamics and mitochondrial dysfunction, highlighting the broad importance of mitochondrial dynamics (Flannery and Trushina, 2019; Pozo Devoto and Falzone, 2017; Van Laar and Berman, 2009; Abati et al., 2024).

Mitofusin-mediated membrane fusion occurs in discrete steps that are dictated by the catalytic cycle (Fig. 1A)(Daumke and Roux, 2017). The mitofusin GTPase domain forms an extensive intermolecular interface when bound to GTP termed the G-G interface (Cao et al., 2017; Qi et al., 2016; Yan et al., 2018; Li et al., 2019). One model for membrane tethering, the first step of fusion, posits that this interface forms across two membranes, or in *trans* (Daumke and Roux, 2017). Atlastin, the fusion DSP required for homotypic ER fusion, mediates membrane tethering via a similar intermolecular GTPase domain interface to that observed in mitofusin, which may be a common feature for fusion DSPs (Bian et al., 2011; Byrnes and Sondermann, 2011). A second model suggests that the mitofusin tethering complex is formed via intermolecular interactions of the C-terminal domain (Dorn, 2019; Koshiba et al., 2004). Importantly, the conformation of the mitofusins when GTP is bound, and therefore the initial tethering step, remains unknown. Progression from tethering to lipid mixing requires that the mitofusin tethering complex dynamically rearranges to pull the opposing membranes together and promote lipid mixing. This likely requires a large conformational change, which is hypothesized to be allosterically coupled to the progressive steps of GTP hydrolysis. Following lipid mixing, disassembly of the fusion complex would include the dissolution of the G-G interface when the catalytic cycle is complete and mitofusin is in the apo state. While this is likely a key feature of disassembly, the conformation of the apo state is not known.

**Figure 1.**
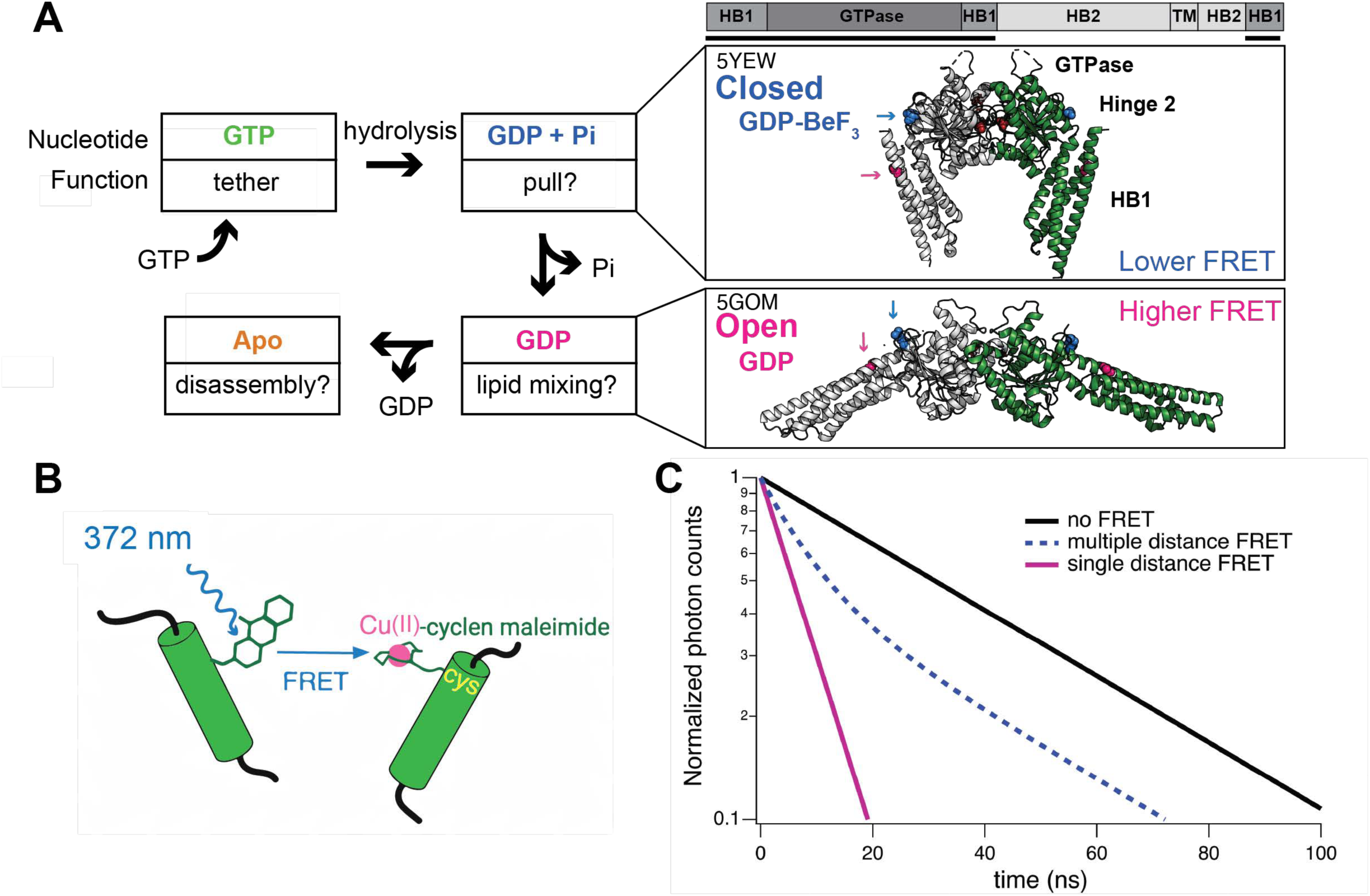
The catalytic cycle drives conformational changes in Mfn1. **(A, left)** Schematic of Mfn1 functional states linked to the nucleotide-bound state throughout the catalytic cycle. At the top left, Mfn1 is bound to GTP, which permits formation of the tethering complex. Hydrolysis of GTP results in GDP + Pi in the nucleotide binding pocket. Using GDP•BeF_3_ to mimic this transition state, a crystal structure of the mini-Mfn1 construct were obtained that revealed the closed state (A, top right [5YEW]). The release of Pi results in the GDP bound state, which could be when lipid mixing occurs. The crystal structure of GDP-mini-Mfn1 showed the open state (A, bottom right [5GOM]). Lastly, GDP is released and no nucleotide is bound. **(A, right)** A schematic denoting the domains found in mitofusin (HB, helical bundle and TM, transmembrane). Black lines below indicate the domains in the crystallization constructs, referred to as mini-mitofusin. The structures are mini-Mfn1 dimers in two distinct nucleotide bound states. As observed in other DSPs, an intermolecular interface between GTPase domains is essential for GTP hydrolysis (G-G interface). The closed conformation was observed with mini-Mfn1 bound to GDP•BeF_3_, mimicking GDP + Pi (5YEW), while the open conformation is GDP-mini-Mfn1 (5GOM). The arrows indicate the areas targeted in our design of the tmFRET donor (pink) and acceptor (blue) positions at L13 and R171, respectively. In this arrangement, the tmFRET pair are further apart in the closed state, which would result in lower FRET efficiency; this pair is closer in the open state, which would result in increased FRET efficiency. **(B)** Schematic of tmFRET with Acd incorporated into the protein and the transition metal ion bound to a cysteine via cyclen-maleimide. Acd is excited at 372 nm and emission is recorded at 446 nm. Fluorescence is quenched by the metal acceptor with an efficiency that is highly dependent on distance. **(C)** Theoretical fluorescence lifetimes of a donor fluorophore in the time-domain showing basis of time-resolved tmFRET. A single exponential donor alone (no FRET), a single distance high-efficiency FRET state (pink) and an example of FRET with multiple donor-acceptor distances are shown (blue dash).

Structural studies have provided some key insights into the mechanisms of mitofusin-mediated fusion (Cao et al., 2017; Qi et al., 2016; Yan et al., 2018; Li et al., 2019). Whereas the structure of full-length mitofusin remains elusive, a minimal GTPase domain construct, mini-Mfn1, containing the GTPase domain and helical bundle 1 (HB1) has provided atomic resolution structures for two nucleotide-bound states (Cao et al., 2017; Qi et al., 2016; Yan et al., 2018). Specifically, the transition state, representing mini-Mfn1 bound to GDP+Pi, is stabilized by salt-bridge contacts between the GTPase domain and HB1 and is referred to as the closed state (Fig. 1A) (Yan et al., 2018). Following the release of Pi, the interaction between the GTPase domain and HB1 is no longer favorable and HB1 moves away from the GTPase domain resulting in a conformation referred to as the open state (Fig. 1A). The role of this large domain rearrangement in the fusion mechanism is unknown. The absence of structural information for wild-type mini-Mfn1 in the GTP-bound state leaves an open question as to whether this is the only large conformation change required for mitofusin-mediated membrane fusion.

To define the conformational dynamics of mitofusin throughout the catalytic cycle, we leveraged our novel transition metal ion fluorescence resonance energy transfer (tmFRET) approach (Zagotta et al., 2021). tmFRET utilizes a donor fluorophore and transition metal ion acceptor site-specifically incorporated in a protein (Fig. 1B). The donor is a fluorescent non-canonical amino acid and the acceptor is a non-fluorescent Cu(II) bound to a metal chelator covalently attached to a solvent-accessible cysteine (Speight et al., 2013). tmFRET can measure distances in the range of 10-50 Å, ideal for analyzing protein conformational changes (Mortensen and Loland, 2020; Gordon et al., 2016). As in classical FRET, the distance between the donor-acceptor pair dictates the FRET efficiency and can be used as a molecular ruler to measure distances between sites on a protein and therefore report different conformational states (Lakowicz, 2006).

To apply the tmFRET system to Mfn1, we utilized mini-Mfn1 (Cao et al., 2017; Qi et al., 2016; Yan et al., 2018). While mini-Mfn1 has a large internal truncation removing HB2 and the TMD, the presence of the C-terminal helix in HB1 is consistent with the domain architecture of the DSP family (Jimah and Hinshaw, 2019; Low et al., 2009). There is also some controversy regarding the topology of Mfn1 (Mattie et al., 2018), but protease protection and C-terminal tagging experiments from our lab support the model that both termini are in the cytoplasm (Engelhart and Hoppins, 2019; Sloat et al., 2019). The mini-Mfn1 construct is an ideal tool since it has robust expression, well developed purification protocols, and a straightforward functional readout (GTPase activity). Furthermore, the available structures of mini-Mfn1 provide a resource to design donor-acceptor pairs with predicted distances within the range of tmFRET.

In this study we used time-resolved tmFRET, quantifying the fluorescent lifetimes of the donor to report FRET efficiency. The fluorescence lifetime, or time between excitation and emission of a photon, is typically on the order of a few nanoseconds (Lakowicz, 2006). Time-resolved FRET has the advantage of recording FRET efficiencies from an entire population of protein in solution as a nanosecond snapshot (Eggan et al., 2024, 2025; Zagotta et al., 2024). This captures distance distributions of the donor-acceptor pair and reveals structural heterogeneity in the protein sample. This heterogeneity can arise from rotameric variations of both the donor and acceptor, flexibility of the protein within any given conformational state, and the presence of multiple conformational states in the sample. In the time-domain, a histogram of the lifetimes of a fluorophore with a single-exponential lifetime appears as a single-exponential decay (linear on a semi-log graph) (Fig. 1C, black). FRET resulting from a single donor-acceptor distance will shorten the lifetime while maintaining the single exponential decay (Fig. 1C, magenta). However, when there are multiple distances, such as from a heterogeneous population of protein, the lifetime will be nonexponential with contributions from all donor-acceptor distances in the sample (Fig. 1C, blue-dashed).

This approach allowed us to capture structural information for each step of the mitofusin catalytic cycle in physiological conditions. Our tmFRET data for mini-Mfn1 in the presence of GDP are in good agreement with the open state crystal structure, establishing that this conformation exists in solution. Interestingly, we found that in the transition state (GDP + Pi) mini-Mfn1 was structurally heterogeneous, with about 60% of the protein in the closed conformation and the rest in the open conformation. Thus, like other ligand-dependent conformational changes we have reported, we can calculate the energetic landscape of the transition state to reveal that the closed state is energetically favorable over the open state (Eggan et al., 2024, 2025). Our approach also allowed structural analysis of both the GTP-bound and apo-mini-Mfn1. We show that the conformation of the GTP-bound mini-Mfn1 was comparable to the open, GDP-bound state. This establishes that both GTP hydrolysis and the release of the free phosphate cause large conformational changes. Finally, a novel conformational state was identified when no nucleotide is bound to mini-Mfn1, probably of relevance to disassembly. In sum, the data presented here represent the first complete analysis of the conformational dynamics of any DSP in solution throughout the catalytic cycle.

## Results

### Generation of the mini-Mfn1 donor-acceptor pair

We sought to use the non-canonical amino acid Acridonylalanine (Acd) as the fluorescence donor and [Cu(cyclenM)]^2+^ as the tmFRET acceptor (Fig. 1B). This donor-acceptor pair has a short R_0_ (the distance which produces 50% FRET efficiency) of 15.3Å, allowing us to monitor only intramolecular FRET, even when mini-Mfn1 is dimerized in the presence of nucleotide. We designed our donor and acceptor sites in the HB1 and GTPase domain, respectively, on either side of the hinge connecting these domains (Fig. 1A). This design would result in the donor and acceptor being further apart in the closed GDP•BeF_3_ state than in the open GDP-bound state resulting in lower and higher FRET efficiencies, respectively.

The metal ion acceptor requires a single solvent accessible cysteine to react with the metal ion chelator. There are six native cysteine residues in the GTPase domain, which were all solvent accessible in solution (data not shown). Therefore, we first engineered a cysteine-less construct (mini-Mfn1-noC) by systematically replacing each cysteine with either alanine or serine, depending on which substitution supported higher expression and robust purification. The GTPase activity of the mini-Mfn1-noC construct was comparable to wild-type protein, indicating that these substitutions did not appreciably alter the catalytic properties (Fig. 2A). To confirm that this construct was functional in the context of full-length Mfn1 in cells, we scored mitochondrial morphology in Mfn1-null mouse embryonic fibroblasts expressing mNeonGreen-Mfn1 variants (Fig. 2B-C). Cells with empty vector possessed a fragmented mitochondrial network, consistent with reduced mitochondrial fusion in the absence of Mfn1. Expression of either murine or human mNeonGreen-Mfn1 restored the reticular mitochondrial network, rescuing the null phenotype. Importantly, we also observed reticular mitochondrial networks in cells expressing a construct of human Mfn1 with the same serine/alanine substitutions in the GTPase domain as generated in mini-Mfn1-noC. This is consistent with this variant supporting mitochondrial fusion activity that is comparable to wild type.

**Figure 2.**
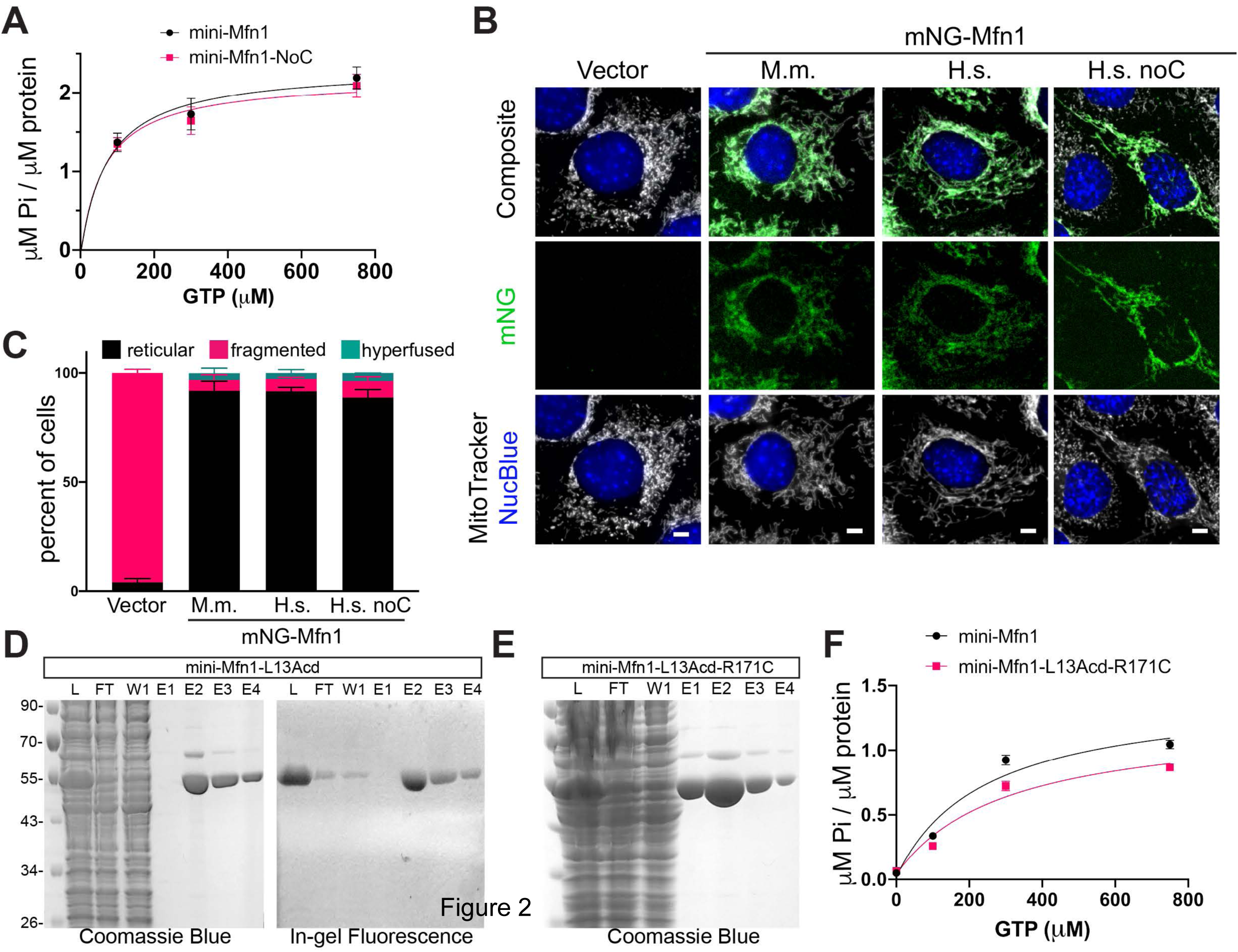
Development of tmFRET donor-acceptor pair on mini-Mfn1. **(A)** GTPase activity of 4.5 μM mini-Mfn1 or mini-Mfn1-NoC measured with a malachite green assay. (p=0.944 [Welch’s unpaired t test]). **(B)** Representative live cell images of Mfn1-null MEF cells with empty vector or expressing the indicated mNeonGreen-Mfn1 construct (mNG-Mfn1)(M.m: *Mus musculus Mfn1*; Hs: *Homo sapiens Mfn1*). Mitochondria were labeled with Mitotracker Red CMXRos and nuclei were labeled with NucBlue; both were visualized by fluorescence microscopy. **(C)** Quantification of mitochondrial morphology for cell lines described in (B). The error bars represent the standard deviation of the five independent and blinded experiments. **(D)** Coomassie gel staining (left) and in-gel fluorescence (right) of indicated fractions from Twin-strep mini-Mfn1-L13Acd purification (Lysate [L], Flowthrough [FT], Wash 1 [W1], Elution 1-4 [E1-4]). Molecular weight markers are indicated on the left (kDa). **(E)** Coomassie gel staining of indicated fractions from Twin-strep mini-Mfn1-L13Acd-R171C purification as in D. **(F)** GTPase activity of 4.5 μM mini-Mfn1-WT or mini-Mfn1-L13Acd-R171C were measured with a malachite green assay. (p=0.7285 [Welch’s unpaired t test]).

We engineered the tmFRET donor site in the first helix of HB1. At this site (L13), we incorporated Acd by amber codon suppression, as previously described (Speight et al., 2013). Coomassie blue staining and in-gel fluorescence of purified TwinStrep-mini-Mfn1 indicate that Acd was successfully incorporated (Fig. 2D). Integration of Acd at only L13 was also supported by mass spectrometry analysis, which revealed no off-target Acd incorporation (Fig. S1). The L13Acd construct lacking the acceptor cysteine (termed donor only) was used as a negative control in all tmFRET experiments. We then introduced the metal-ion acceptor site at R171, which is predicted to be within tmFRET range from L13 in both the open and closed states (13.8 Å and 24.0 Å, respectively). This construct (mini-Mfn1-L13Acd-R171C) also expressed robustly (Fig. 2E). The GTPase activity of mini-Mfn1-L13Acd-R171C was also comparable to wild-type controls (Fig. 2F), indicating that these mutations did not alter catalytic activity.

### Time-resolved tmFRET reveals distance distributions of mini-Mfn1 in GDP- and transition-bound states

With our functional mini-Mfn1-L13Acd-R171C protein, we next sought to measure the donor-acceptor distances, starting with the two ligands previously used for X-ray crystallography with mini-Mfn1-WT bound to GDP or GDP•BeF_3_, which mimics GDP + Pi (Fig. 3A, D). We measured fluorescence lifetimes in the form of time-correlated single photon counting (TCSPC) decays. We first established the lifetime of the donor-only mini-Mfn1-L13Acd-noC in the presence of saturating GDP (brown trace, Fig. 3B), which produced decays similar to that of Acd-incorporated proteins reported previously (Eggan et al., 2025). We then measured the lifetime of Acd in the donor only mini-Mfn1-L13Acd-noC protein construct in the presence of [Cu(cyclenM)]^2+^, which is not expected to react with this cysteine-less protein (violet trace, Fig. 3B). As expected, this lifetime was very similar to that obtained in the absence of [Cu(cyclenM)]^2+^. Similar results were obtained for the donor-only mini-Mfn1-L13Acd-noC in the presence of saturating GDP•BeF_3_ (Fig. 3E). This demonstrates that unreacted [Cu(cyclenM)]^2+^ did not quench Acd in the protein and did not bind to off-target sites near the Acd.

**Figure 3.**
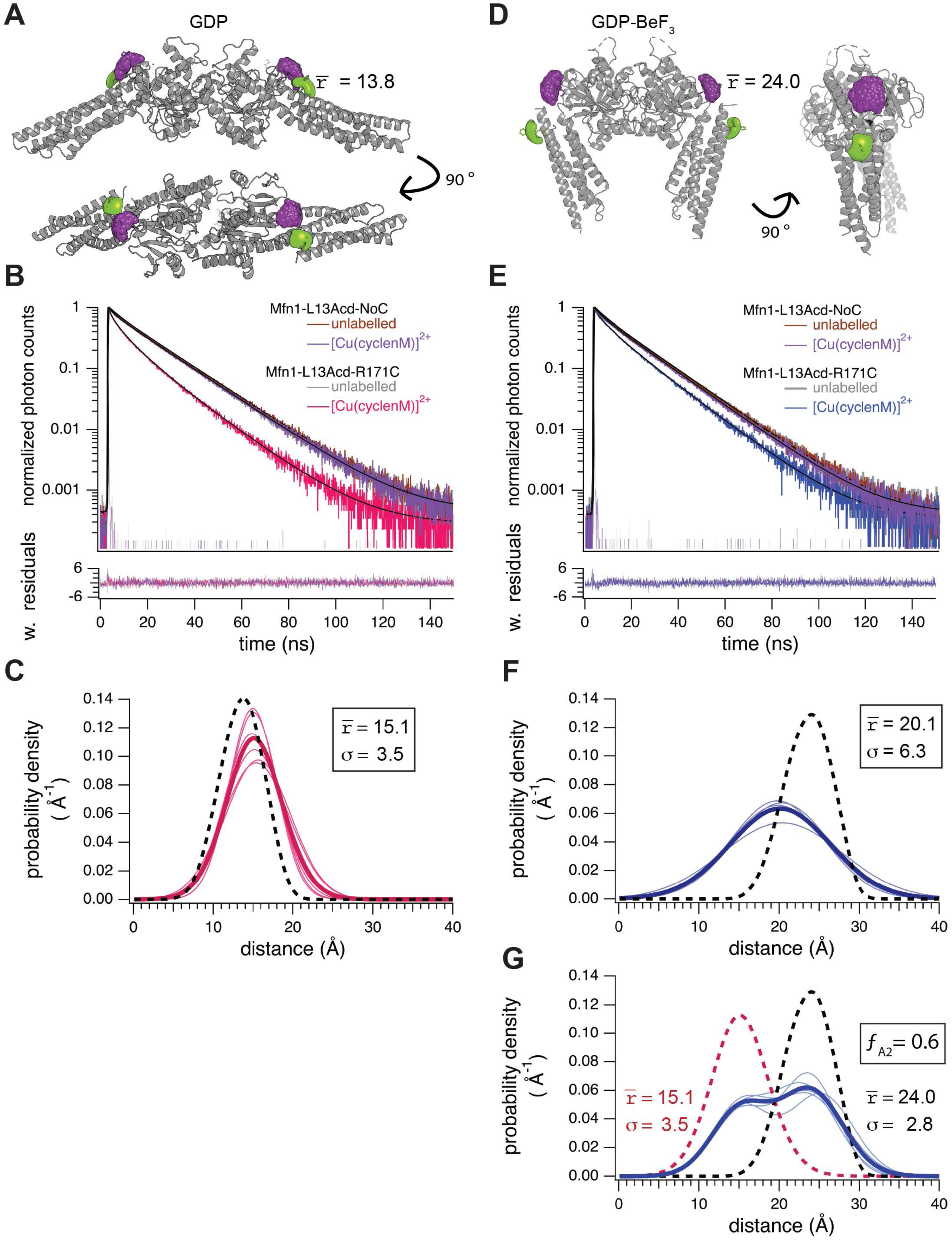
Steady-state lifetime tmFRET of mini-Mfn1 in the GDP-bound and transition states. (**A**) Cartoon structure of mini-Mfn1 in the GDP-bound state with the predicted rotameter states of L13Acd (green) and R171[Cu(cyclenM)]^2+^ (purple) depicted as a cloud. The average predicted donor-acceptor distance is 13.7 Å. (**B**) Representative TCSPC data from saturating GDP conditions with either donor only (Mfn1-L13Acd-NoC) before (brown) and after (violet) the addition of [Cu(cyclenM)]^2+^ or from mini-Mfn1-L13Acd-R171C before (grey) and after (magenta) the addition of [Cu(cyclenM)]^2+^. Weighted residuals are shown below using the same colors as used for the raw data above. (**C**) Spaghetti plot of distributions from individual experiments with GDP with average of all experiments in bold, and the distribution predicted by chiLife in the dashed black curve. (**D**) Cartoon structure of mini-Mfn1 in the transition state (GDP•BeF_3_) with the predicted rotameter states of L13Acd (green) and R171[Cu(cyclenM)]^2+^ (purple) depicted as a cloud. The average predicted distance is 24.4 Å. (**E**) Representative TCSPC data from saturating conditions of GDP•BeF_3_ with either donor only (Mfn1-L13Acd-NoC) before (brown) and after (violet) the addition of [Cu(cyclenM)]^2+^ or Mfn1-L13Acd-R171C before (grey) and after (blue) the addition of [Cu(cyclenM)]^2+^. Weighted residuals are shown below using the same colors as used for the raw data above. (**F**) Spaghetti plot of distributions from individual experiments with GDP•BeF_3_ fit to a single Gaussian with the average of all experiments in bold, and the distribution predicted by chiLife in the dashed black curve. The reduced chi-square values associated with the fit of this model for each of the individual replicates are as follows: rep1=0.67, rep 2=0.66, rep 3=0.64, rep 4=0.69, rep 5=0.74, rep 6=0.78. (**G**) Spaghetti plot of distributions from individual experiments with GDP•BeF_3_ fit to a dual Gaussian with the average in bold (blue). The GDP•BeF_3_-bound state distribution predicted by chiLife (dashed black curve) and the experimental average GDP distribution (dashed magenta curve) are overlayed. The fraction in the closed state (ƒ_A2_) is 0.6. The reduced chi-square values associated with the fit of this model for each of the individual replicates are as follows: rep1=0.67, rep 2=0.67, rep 3=0.65, rep 4=0.7, rep 5=0.75, rep 6=0.84.

We next measured the lifetime of the complete tmFRET construct, mini-Mfn1-L13Acd-R171C-[Cu(cyclenM)]^2+^, in the presence of GDP. The acceptor decreased the overall lifetime of Acd, consistent with FRET (magenta, Fig. 3B). We fit the lifetimes with a model that assumes a Gaussian distribution of donor-acceptor distances. The average distance (r̅) and standard deviations (σ) from multiple experiments are shown in a spaghetti plot in Figure 3C with a mean distance r̅ = 15.1 Å and standard deviation σ = 3.5 Å. To generate a predicted distance distribution for comparison to our experimental data, we used chiLife. chiLife is a software package that models rotameric ensembles of the donor and acceptor on a stationary backbone to generate a predicted distance distribution for the donor-acceptor pair (Fig. 3A)(Tessmer and Stoll, 2023). The distance distribution predicted for GDP-bound Mfn1 (PDB:5GOM) by chiLife is overlayed on the spaghetti plot and has an average r̅ = 13.8 Å and σ = 2.7 Å (dotted line). This represents a strong agreement between the prediction and our experimentally determined distribution for both the average distance and the standard deviation (Fig. 3C). The small differences from the chiLife predictions are likely due to the differences in energetic and geometric constraints that exist in the crystallization conditions but are absent in solution.

The same experiments were performed with mini-Mfn1-L13Acd-R171C-[Cu(cyclenM)]^2+^ in the presence of saturating GDP•BeF_3_ (blue, Fig. 3E). Compared to GDP conditions, we measured a smaller change after labeling with [Cu(cyclenM)]^2+^ in the GDP•BeF_3_ condition. This is consistent with the prediction that the donor-acceptor distance is greater in the closed, GDP•BeF_3_-bound state compared to the GDP-bound state (Fig. 3A, D). The distance distributions fit to a single state for GDP•BeF_3_-mini-Mfn1 are summarized as spaghetti plots in Figure 3F, with an average r̅ = 20.1 Å and σ = 6.3 Å (Tessmer and Stoll, 2023). The distance distribution predicted for GDP•BeF_3_-bound Mfn1 (PDB:5YEW) by chiLife is overlayed on the spaghetti plot and has an average r̅ = 24.0 Å and σ = 2.8 Å (black-dashed). Compared to the chiLife prediction, the experimentally determined r̅ value is smaller and the σ is larger, resulting in a shifted and wider Gaussian curve. This model would position HB1 and the GTPase domain further apart than observed in the GDP•BeF_3_ crystal structure with greater conformational heterogeneity than predicted based on rotomeric heterogeneity alone. In fact, this distance between HB1 and the GTPase domain is unlikely to permit the formation of salt bridges. These data are not in strong agreement with the predictions based on the crystal structure, prompting us to consider additional interpretations of the data.

Given the large σ obtained from our fits to the lifetime data, we also considered the possibility that the data represents the presence of two populations - one in the closed state and a second in the open state. This would be consistent with a model where GDP•BeF_3_ alters the energetics of mini-Mfn1, making the closed state more favorable than when bound to GDP. To test whether our data support a distribution of the protein among two conformational states, we fit the data with a model that specified two Gaussian donor-acceptor distance distributions, our experimental GDP-mini-Mfn1 and the chiLife prediction of the closed state, based on the X-ray crystal structure. The fraction in the closed state (ƒ_A2_), and therefore the distribution among the two Gaussians, was allowed to vary. In the context of this two-state model, our data indicate that about 60% of mini-Mfn1 is in the closed conformation in the presence of GDP•BeF_3_ and the remaining fraction is in the open state (Fig. 3G). We obtained similar chi-squared values from fits with both single- and two-state models to our data indicating that they are statistically equivalent.

### A known functional variant supports the two-state model of GDP•BeF_3_ bound mini-Mfn1

We used a known functional variant, R73Q, to test our hypothesis that a two-state model for GDP•BeF_3_ can best explain our data. The GDP•BeF_3_ crystal structure identified salt bridges between R73 in HB1 and D200 and D198 in the GTPase domain (Yan et al., 2018). The importance of the salt bridges was demonstrated by characterizing variants with charge reversal substitutions in either domain. These variants possessed defects in dimerization of mini-Mfn1 *in vitro* and in mediating mitochondrial fusion in cells. In addition, a common CMT2A-related R→Q variant was assessed by bulk FRET experiments with purified mini-Mfn1-R73Q to show that this variant does not readily access the closed conformation, although dimerization was unchanged (Yan et al., 2018). These data all support a model in which these salt bridges stabilize the closed state when GDP•BeF_3_ is bound; thus, in a two-state model, disruption of the salt bridges is expected to shift the equilibrium toward the open state.

We created the R73Q substitution in our mini-Mfn1-L13Acd-R171C construct to assess the predictions derived from the two-state model. If GDP•BeF_3_-mini-Mfn1 is in equilibrium between the open and closed states, we expect that the R73Q variant will primarily exist in the open state; however, if GDP•BeF_3_-mini-Mfn1 exists in a single state with 20 Å distance between the donor and acceptor, we expect that the absence of the salt bridges would be less impactful and the substitution of R73Q might induce a third novel conformational state with an r̅ between the GDP and GDP•BeF_3_-bound values.

We first measured the lifetimes of mini-Mfn1-L13Acd-R73Q-R171C before and after adding [Cu(cyclenM)]^2+^ in the presence of GDP (Fig. 4A). A summary of these experiments is shown in the spaghetti plot in Figure 4B with the average experimental GDP-mini-Mfn1 distribution plotted in the dashed line. This revealed an average distance distribution of r̅ = 15.2 Å (σ = 4.6 Å) for the mini-Mfn1-L13Acd-R73Q-R171C, comparable to the wild-type protein. This suggests the salt bridge has little or no effect on the structure or energetics of the GDP-bound state, as expected.

**Figure 4.**
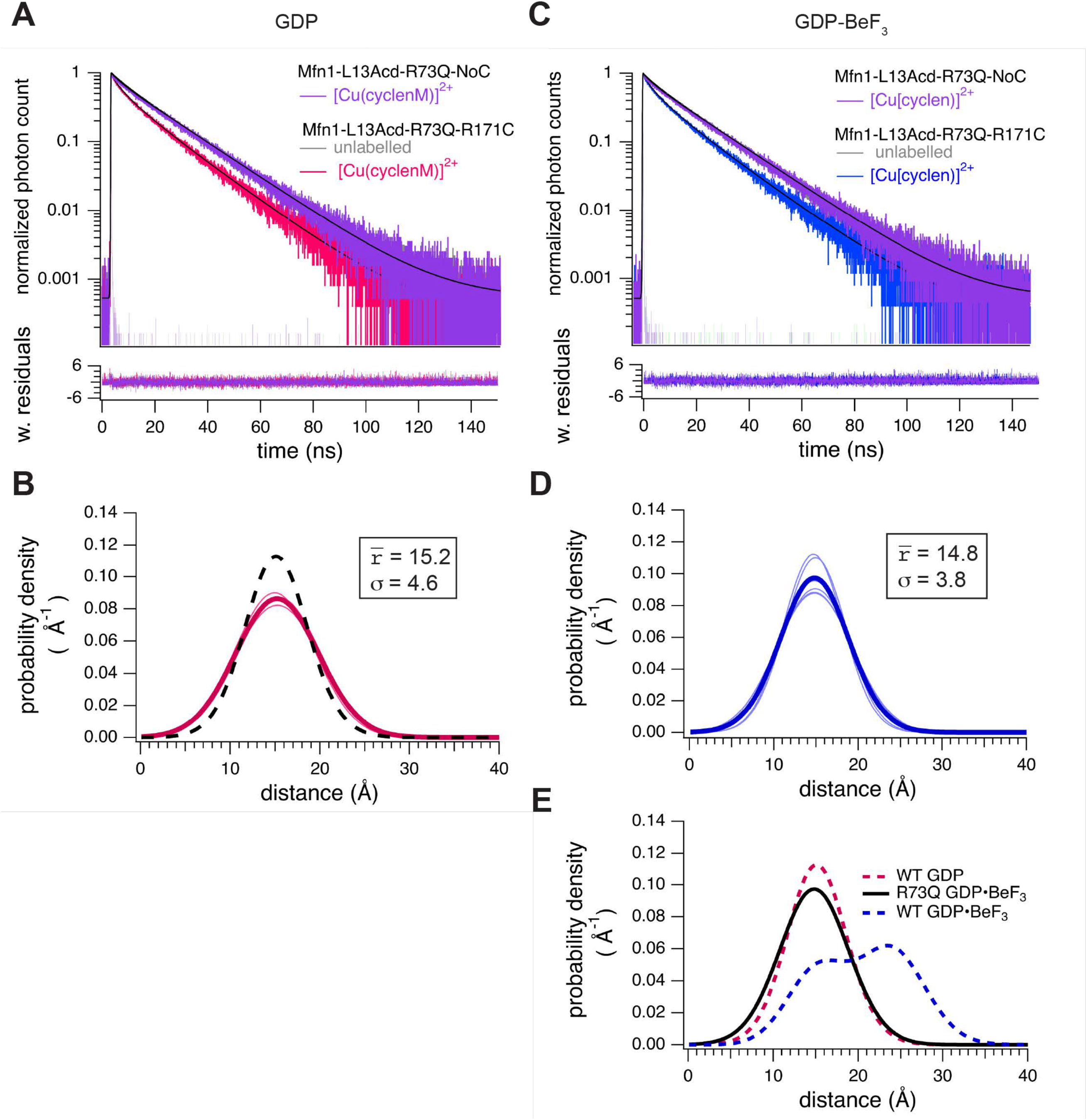
Steady-state lifetime tmFRET of mini-Mfn1-R73Q in the transition- and GDP-bound states. (**A**) Representative TCSPC data from saturating GDP conditions with either donor only (Mfn1-L13Acd-R73Q-NoC) after (violet) the addition of [Cu(cyclenM)]^2+^ or from mini-Mfn1-L13Acd-R73Q-R171C before (grey) and after (magenta) the addition of [Cu(cyclenM)]^2+^. Weighted residuals are shown below using the same colors as used for the raw data above. (**B**) Spaghetti plot of distributions from individual experiments with GDP. Average is in bold, and wild-type distribution shown in the dashed black curve. (**C**) Representative TCSPC data from saturating conditions of GDP•BeF_3_ with either donor only (Mfn1-L13Acd-R73Q-NoC) after (violet) the addition of [Cu(cyclenM)]^2+^ or Mfn1-L13Acd-R73Q-R171C before (grey) and after (blue) the addition of [Cu(cyclenM)]^2+^. Weighted residuals are shown below using the same colors as used for the raw data above. (**D**) Spaghetti plot of distributions from individual experiments with GDP•BeF_3_ with the average in bold (**E**) Plot of average distributions from Mfn1-L13Acd-R73Q-R171C in GDP•BeF_3_ (black) compared to wildtype in GDP (magenta) and GDP•BeF_3_ (blue). (F) Spaghetti plot of distributions from individual experiments with GDP•BeF3 fit to a single Gaussian with the average of all experiments in bold, and the distribution predicted by chiLife in the dashed black curve. (G) Spaghetti plot of distributions from individual experiments with GDP•BeF_3_ fit to a dual Gaussian with the average in bold (blue). The GDP•BeF3-bound state distribution predicted by chiLife (dashed black curve) and the experimental average GDP distribution dashed magenta curve) are overlayed.

We then measured the lifetime of mini-Mfn1-L13Acd-R73Q-R171C before and after adding [Cu(cyclenM)]^2+^ in the presence of GDP•BeF_3_ (Fig. 4C). A summary of these experiments is shown in the spaghetti plot in Figure 4D with an average distance distribution of r̅ = 14.8 Å (σ = 3.8 Å). This distance is much shorter, and the Gaussian is narrower than the wild-type protein in the presence of GDP•BeF_3_ (Fig. 4E). In fact, this Gaussian is very similar to wild-type protein in the open state (GDP bound). These data suggest that the energetics of the Mfn1-R73Q variant in the presence GDP•BeF_3_ strongly favor the open state, with little-to-no probability of being in the closed state. Therefore, our analysis supports the two-state model for the GDP•BeF_3_ condition as the distance distribution of mini-Mfn1-R73Q matches the open state with a small standard deviation.

Because time-resolved tmFRET data capture a nanosecond snapshot of the conformational distribution, they encode information about the free energy difference between states and how this free energy difference changes in response to ligand. In this case, the proportion of molecules in each state can be directly converted into a change in the Gibbs free energy (ΔG) for the transition of HB1 relative to the GTPase domain from open to closed. For wild-type mini-Mfn1, we calculate a free energy difference between closed and open states with GDP•BeF_3_ bound of ΔG= −0.21 kcal/mol, indicating that closed state is energetically more favorable. This represents the only known nucleotide condition where the closed state is favored. Given that mini-Mfn1-R73Q is believed to be primarily in the open conformation, the inter-domain salt bridges contribute significantly to this energetic effect.

### Non-hydrolyzable GTP analogs induce the open conformation of mini-Mfn1

We next examined the GTP-bound state of mini-Mfn1. Cao *et al*. obtained the structure of a catalytic dead, monomeric mini-Mfn1-T109A bound to GTP in the open state, comparable to GDP-mini-Mfn1; however, the structure of wild-type Mfn1 bound to GTP is not known. We utilized two different non-hydrolyzable GTP analogs, since it has been previously shown that each non-hydrolyzable GTP analog produces slight functional differences in DSPs (Byrnes and Sondermann, 2011; Shi et al., 2024). We measured the lifetimes of mini-Mfn1-L13Acd-R171C with either GTPγS or GMP-PNP before and after adding [Cu(cyclenM)]^2+^ (Fig. 5A & B). Our analysis yielded average distances of r̅ = 14.6 Å (σ = 3.7 Å) when incubated with GTPγS and r̅ = 15.5 Å (σ = 3.6 Å) with GMP-PNP (Fig. 5C & D). Comparing these distributions to the distributions we determined from incubation with GDP reveals minimal differences in the donor-acceptor distance distributions (Fig. 5E). Based on these data we conclude that the GTP-bound state is comparable to the open, GDP-bound conformation.

**Figure 5.**
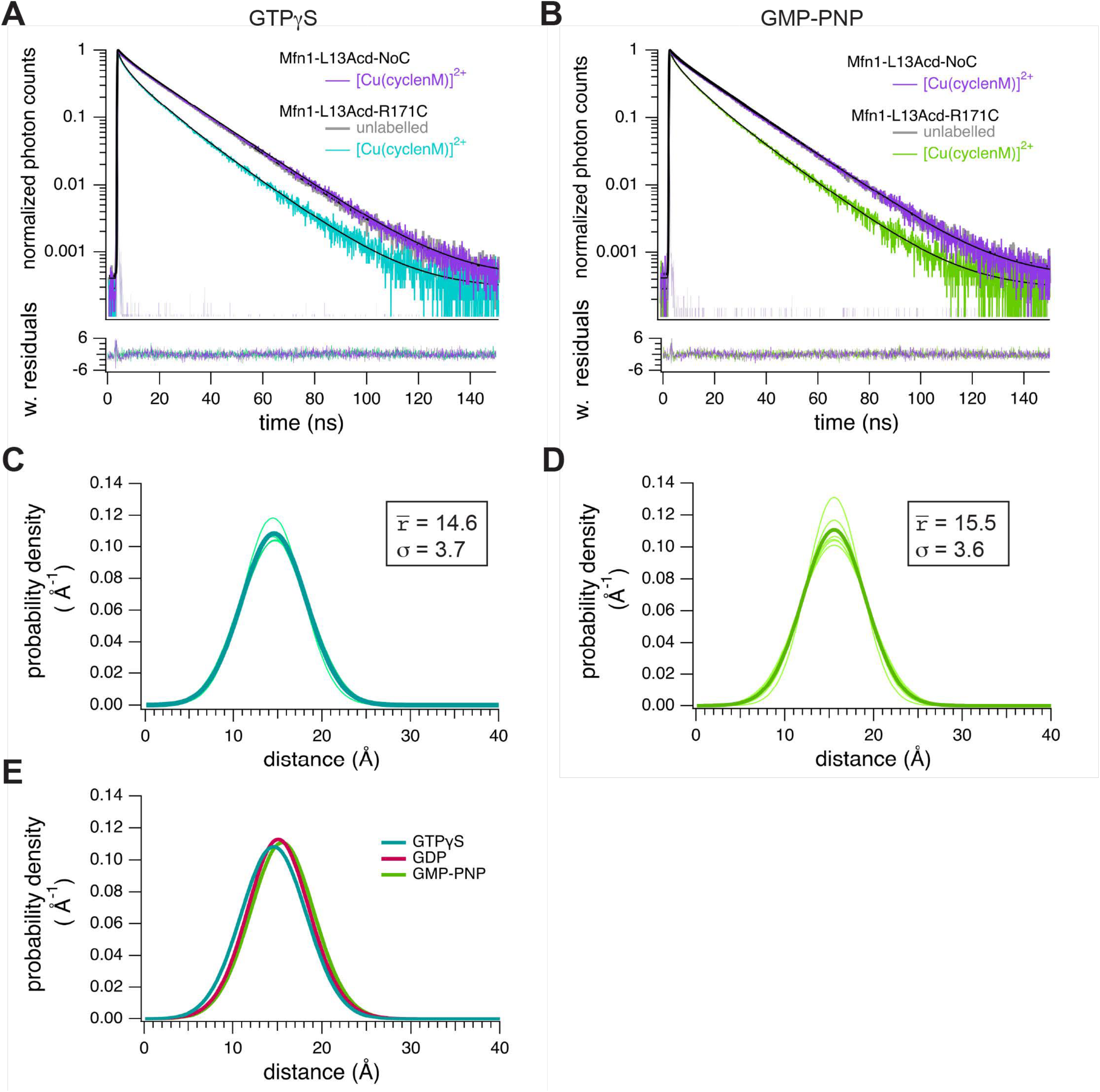
Steady-state lifetime tmFRET of mini-Mfn1 with GTPyS or GMP-PNP bound. (**A)** Representative TCSPC data from saturating GTPyS conditions with either donor only (Mfn1-L13Acd-NoC) after (violet) the addition of [Cu(cyclenM)]^2+^ or from mini-Mfn1-L13Acd-R171C before (grey) and after (cyan) the addition of [Cu(cyclenM)]^2+^. Weighted residuals are shown below using the same colors as used for the raw data above. (**B**) Spaghetti plot of distributions from individual experiments with GTPyS. Average is in displayed in bold. (**C**) Representative TCSPC data from saturating GMP-PNP conditions with either donor only (Mfn1-L13Acd-NoC) after (violet) the addition of [Cu(cyclenM)]^2+^ or from mini-Mfn1-L13Acd-R171C before (grey) and after (green) the addition of [Cu(cyclenM)]^2+^. Weighted residuals are shown below using the same colors as used for the raw data above. (**D**) Spaghetti plot of distributions from individual experiments with GMP-PNP. Average wave is displayed in bold. (**E**) Plot of average distributions for GTPyS (cyan), GDP (magenta), and GMP-PNP (green).

These data capture each nucleotide-bound state in the mini-Mfn1 catalytic cycle. The simplest interpretation is that the closed conformation is favored specifically after GTP hydrolysis, when GDP + Pi occupies the nucleotide pocket. Together, these data support a model in which HB1 can close upon GTP hydrolysis and reopens following release of Pi within a single catalytic cycle.

### Time resolved tmFRET reveals a novel conformation when mini-Mfn1 is not bound to nucleotide

We next assessed the conformation of mini-Mfn1 in the absence of nucleotide, which occurs before and after GTP hydrolysis. The previously solved crystal structure of apo-mini-Mfn1 closely resembled the open GDP-mini-Mfn1. Notably, in the absence of nucleotide, the GTPase domain was not competent for G-G interface formation because amino acid side chains required for salt bridge interactions were inaccessible (Cao et al., 2017). As this is the only intermolecular interface in the mini-Mfn1 construct, the apo-mini-Mfn1 is the only state that must exist exclusively as a monomer.

We measured the lifetime of mini-Mfn1-L13Acd-R171C with no nucleotide before and after adding [Cu(cyclenM)]^2+^ (Fig. 6A). A summary of these experiments is shown in the spaghetti plot in Figure 6B. Under these conditions, we calculated an average distance between the donor and acceptor of r̅ = 16.3 Å and σ = 4.4 Å. The distance distribution predicted for nucleotide free mini-Mfn1 (PDB:5GO4) by chiLife is overlayed on the spaghetti plot and has an average r̅ = 14.8 Å and σ = 2.7 Å (black-dashed). While both GTP-bound and GDP-bound exhibit average distances of ∼15Å between the FRET pair, the apo state consistently results in a distance slightly greater than 16Å, suggesting this a unique conformational state that exists when mini-Mfn1 is not constrained by the energetic requirements of crystal packing. Indeed, the difference in average distance for GDP-mini-Mfn1 and apo mini-Mfn1 (r̅) is statistically significant (Fig. 6C & 6D). Furthermore, the σ is significantly larger for apo-mini-Mfn1 compared to the GDP-bound state, indicating increased flexibility of this conformational state (Fig. 6C & 6E).

**Figure 6.**
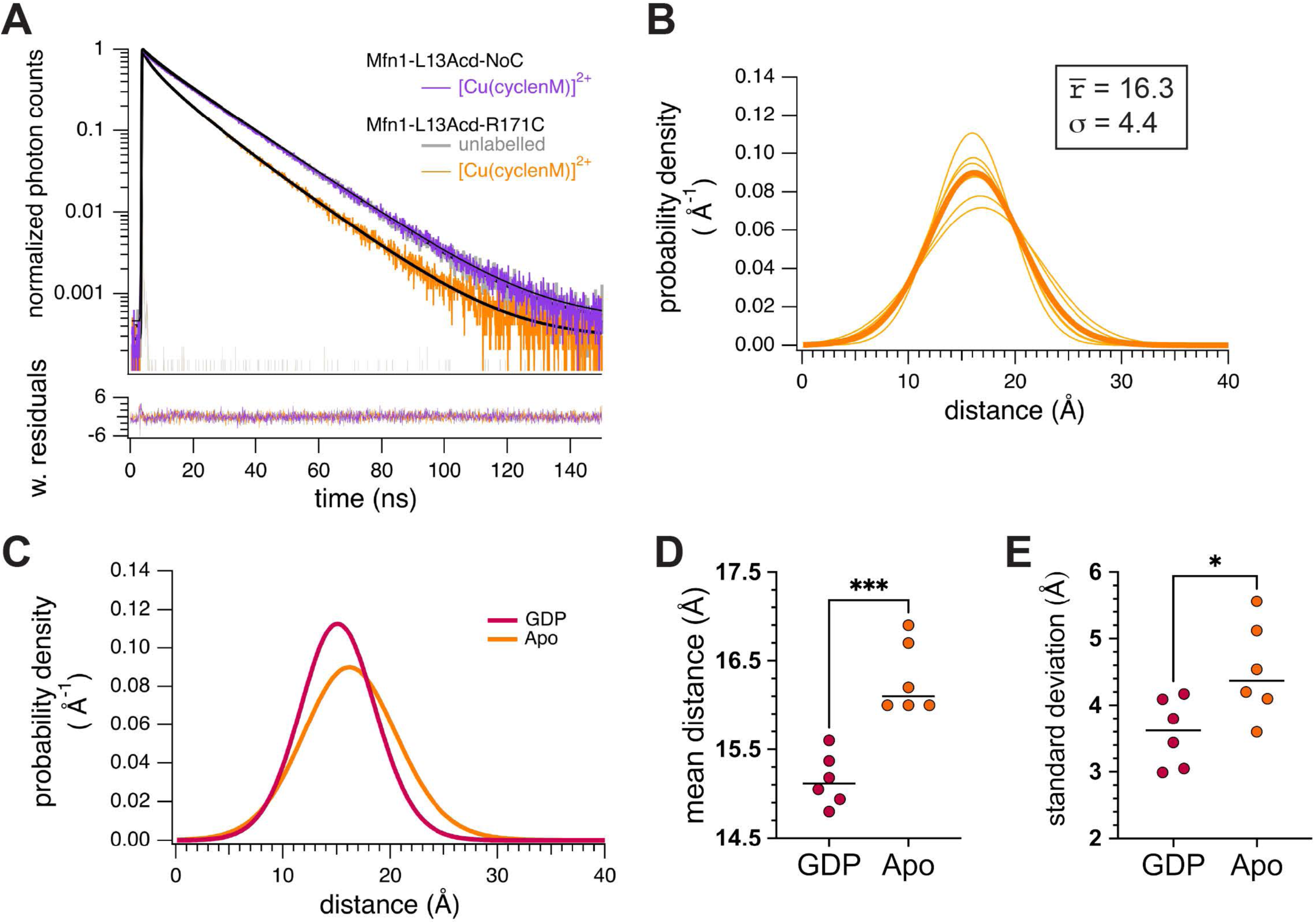
Steady-state lifetime tmFRET of mini-Mfn1 in the apo state. (**A**) Representative TCSPC data with either donor only (Mfn1-L13Acd-NoC) after (violet) the addition of [Cu(cyclenM)]^2+^ or from mini-Mfn1-L13Acd-R171C before (grey) and after (orange) the addition of [Cu(cyclenM)]^2+^. Weighted residuals are shown below using the same colors as used for the raw data above. (**B**) Spaghetti plot of distributions from individual experiments with no nucleotide bound fit to a single Gaussian distribution. The average distribution is displayed in bold. The reduced chi-square values associated with the fit of this model for each of the individual replicates are as follows: rep1=0.67, rep 2=0.61, rep 3=0.71, rep 4=0.70, rep 5=0.69, rep 6=0.70. (**C**) Plot of average distributions for GDP (magenta) and apo (orange) **(D)** Mean distances for individual experiments are compared either in the presence of GDP or in the absence of nucleotide (Apo) where *** P<0.01 and * P<0.05 (Welch’s unpaired t test). **(E)** Sigma values for individual experiments are compared either in the presence of GDP or in the absence of nucleotide (Apo) where *** P<0.01 and * P<0.05 (Welch’s unpaired t test).

As described above, a large sigma can indicate that there are two distinct conformational states present in that condition. The sigma for the single-state model of apo-mini-Mfn1 (4.4 Å) is not as large as observed for the GDP•BeF_3_-bound state (6.3 Å); nonetheless we fit the same data to the two-state model used for GDP•BeF_3_ (Fig. 3G). The model estimates that a minor fraction of the total protein would be in the closed state (ƒ_A2_ = 0.26)(Fig. S2). Analysis of the fit to each model resulted in similar reduced chi-square values, indicating that they are statistically equivalent.

To further probe the apo state, we utilized a functional variant carrying an alanine substitution at R238, the side chain of which becomes solvent-facing following GTP binding, forming an essential salt bridge across the G-G interface. The Mfn1-R238A variant is catalytically dead, unable to dimerize via the G-G interface and thus unable to hydrolyze GTP (Cao et al., 2017). Therefore, mini-Mfn1-R238A is predicted to be largely functionally equivalent to the apo state and would be refractory to any conformational changes that require the G-G interface or GTP hydrolysis. Consistent with this, Mfn1-R238A recapitulated the apo state in the presence of each ligand, with distance averages of ∼16 Å or more and σ values ≥ 4.6 (Fig. S3). As shown for apo-mini-Mfn1, the R238A data also fit a two-state model with an ƒ_A2_ of 0.2 and 0.3 in the apo and GDP•BeF_3_ conditions, respectively (Fig. S3).

These data reveal a conformational state of mini-Mfn1 distinct from both the open and closed states characterized in nucleotide-bound conditions. The most parsimonious interpretation is the single-state model where the donor and acceptor pair further apart than the GTP- and GDP-bound open state. Furthermore, the larger sigma than predicted by chiLife suggests that this conformation is flexible. Therefore, GTP binding would favor movement of HB1 upwards and stabilize the open state. Given that the apo state occurs at the end of the catalytic cycle, this unique conformational state may play a role in the disassembly of the post-fusion complex.

## Discussion

Here, we established a time-resolved tmFRET approach to measure the conformational dynamics of mini-Mfn1. Time-resolved tmFRET reliably provides valuable structural information in the form of average distances of each conformational state, heterogeneity of the distance for each conformation, and the relative proportion of different conformations (energetics) in a population. This allowed us to acquire structural information for every step of the catalytic cycle, and insight into the energetics of the large conformational change regulated by GTP hydrolysis, neither of which was possible with previous approaches.

Mitochondrial fusion is initiated by the interaction of two mitofusin complexes on distinct mitochondrial membranes, representing the formation of a tethering complex. To pull the membranes together and promote lipid mixing, this tethering complex must dynamically rearrange through conformational changes that are allosterically regulated by the catalytic cycle. We have obtained several novel insights into the GTP-dependent allosteric regulation of Mfn1 (summarized in Fig. 7). Our data revealed the conformational state of wild type Mfn1 in the GTP-bound state (Figure 5). GTP-binding is associated with the formation of the tethering complex, and our data indicate that Hinge 2 is in the open state in this complex. We also obtained novel insights into the catalytic step following GTP hydrolysis, where Mfn1 was previously reported to be in a closed conformation with HB1 interacting with the GTPase domain. The time-resolved tmFRET analysis revealed that the transition state results in an equilibrium between the closed and open states (Fig. 3G). The energetic calculations demonstrate that about 60% of the population is in the closed state. This conclusion was supported by our analysis of mini-Mfn1-R73Q, which alters the salt bridge network between the GTPase and HB1 domains, and was only in the open state (Fig. 4). Previous bulk FRET approaches to study the open-to-closed conformational change utilized fluorescence intensity to calculate FRET (Yan et al., 2018). In that case, the change in FRET intensity could only identify the time-average conformation of the population, but not the proportion of mini-Mfn1 in each state. This indicates that the energetic landscape of mitofusin-mediated fusion is more nuanced than other membrane fusion machines, such as SNAREs or viral membrane fusion proteins, which are modelled as largely irreversible conformational changes driving membrane fusion. Not only is the transition state in equilibrium between open and closed, but the conformational change that results from GTP hydrolysis is reversed by release of Pi. This is supported by our analysis of the GDP-bound state, which revealed a calculated distance in solution that is remarkably comparable to chiLife predictions (Fig. 3). Together with our GTP-bound conformational data, we propose a model of mitofusin-mediated fusion in which the open GDP-bound state represents the second conformational change of a single catalytic cycle, back to a relatively stable open state. We speculate that this movement of the GTPase domain away from HB1 may allow for a large conformational change in the position of HB2, to promote lipid mixing. Finally, this approach also enabled structural analysis of Mfn1 in the absence of nucleotide. These data are most consistent with this being a novel position of HB1 relative to the GTPase domain with more conformational heterogeneity than observed in the GTP- and GDP-bound states (Fig. 6). This indicates that GTP-binding alters the conformational dynamics to favor the open state. The distinct apo state is likely significant for the disassembly of the fusion complex and therefore a key step in the mechanism.

**Figure 7.**
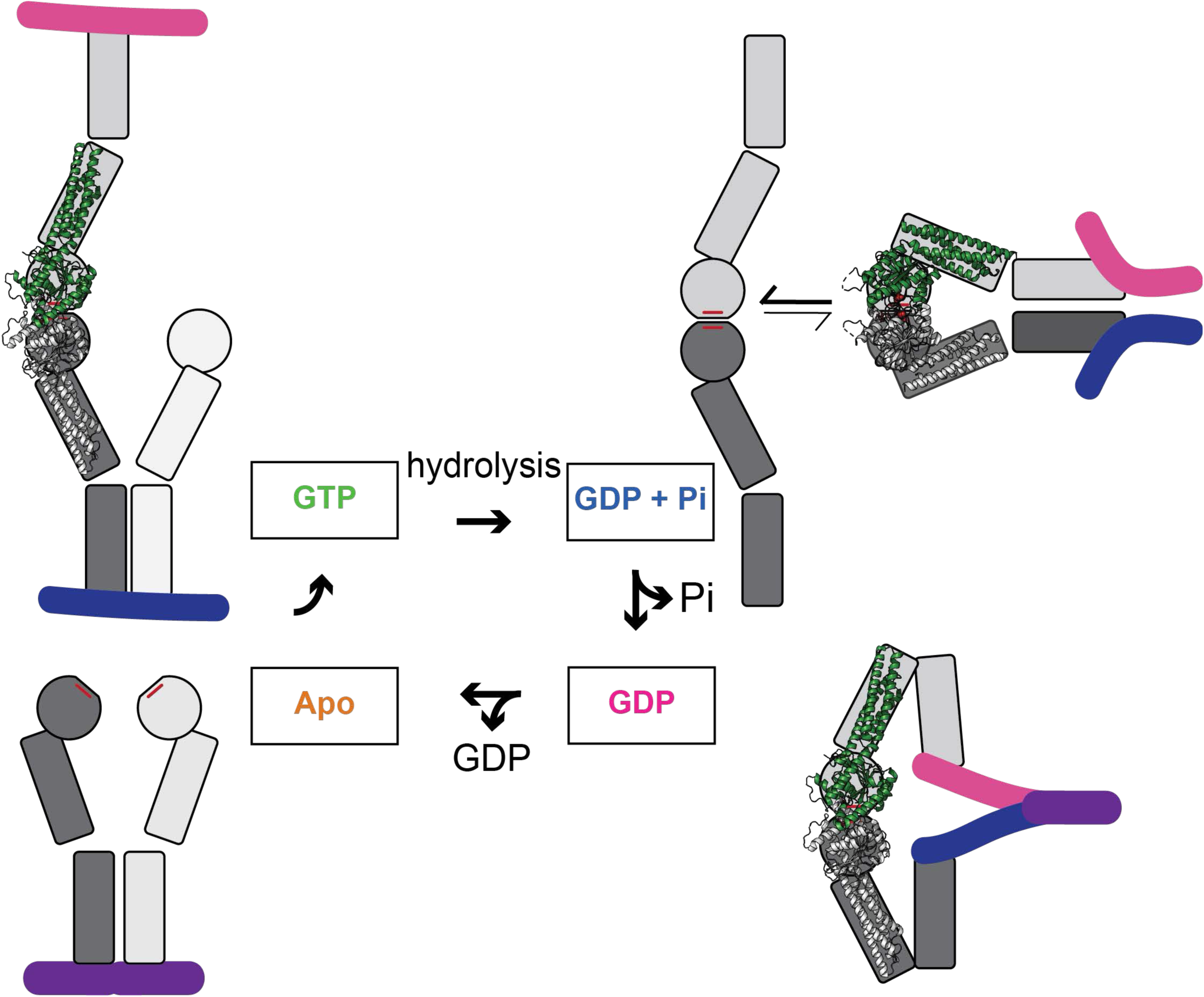
A model for the conformational changes driven by the catalytic cycle of Mfn1. To mediate mitochondrial outer membrane fusion, mitofusins establish a physical connection between two mitochondria via formation of tethering complexes. We have previously shown that full-length Mfn1 in the mitochondrial outer membrane oligomerizes in the same membrane (in *cis*) as well as across two membranes (in *trans*). Therefore, we depict *cis* dimers in the first and last panel but remove this for simplicity in the intermediate panels. Our data demonstrate that the GTPase and HB1 domains are in the open state for GTP-Mfn1, which supports membrane tethering. We depict HB2 as grey boxes and the membrane as pink and blue lines. The tethering complex must dynamically re-arrange to pull the two membranes together and promote lipid mixing. These conformational changes are allosterically controlled by the catalytic cycle. Following GTP hydrolysis, GDP+Pi is found in the binding pocket, and our data indicate an equilibrium between the open and closed states, with the closed state favored. We predict that this conformational change of HB1 relative to the GTPase domain will pull the two membranes together and could result in new intermolecular interface(s) involving HB2. Release of Pi generates GDP-Mfn1, which favors the open state. We present a model where a large conformational change of the HB2 domain relative to HB1 generates a membrane structure that favors lipid mixing. When GDP is released and Mfn1 is in the apo state, our data indicate that the GTPase domain and HB1 are in unique positions compared to the other states.

Given that mini-Mfn1 is in equilibrium between the closed and open state when bound to GDP + Pi, we predict that this could be an important point of regulation in mitochondrial fusion efficiency. For example, the energetics could be altered to favor the closed state, which we predict would favor membrane fusion. This may be observed in a heterotypic Mfn1-Mfn2 tethering complex, which is known to be the superior fusion machine (Hoppins et al., 2011). This could also be affected by cytosolic regulators that promote fusion, such as Bax. In addition, the presence of HB2 and/or the membrane load could alter the energetic landscape to favor or disfavor the closed state. In our model presented in Figure 7, we speculate that intermolecular interactions between HB2 domains could favor the closed state conformation of HB1 relative to the GTPase domain.

One key limitation to our work is the use of the truncated mini-Mfn1, which is missing HB2 and is not embedded in a membrane. Both are essential aspects of the mechanism of mitofusin-mediated fusion and could alter the dynamics of Hinge 2 throughout hydrolysis. Indeed, allosterically regulated conformational changes are predicted to propagate throughout the entire protein. Nonetheless, this system has revealed fundamental aspects of the conformational energetics of Mfn1, exemplified by the fact that we identified two conformational states that had not previously been explored. We propose a model for the role of HB2 in mitochondrial outer membrane fusion (Figure 7). In this model, HB2 would play a role in stabilizing the closed state of Hinge 1, which would also be associated with a large change in the proximity of the two membranes relative to the initial tethered state. The release of Pi reverts the positions of the GTPase and HB1 domains back to a position observed in the GTP-bound state. We speculate that this could make room for a large conformational change by HB2 that would promote lipid mixing (Figure 7, bottom left).

Future work will expand to applying this system to the other mitofusin paralog, Mfn2, as well as to full-length mitofusin in membranes. Importantly, Mfn1 and Mfn2 that have fundamental differences in catalytic function, with mini-Mfn2 having an 8-fold lower hydrolysis rate compared to mini-Mfn1 (Li et al., 2019). Furthermore, past work has shown that mitochondrial fusion is most efficient when the different paralogs are on opposing membranes (Hoppins et al., 2011). The heterotypic fusion complex could result in altered conformational energetics of each paralog to increase fusion efficiency. Alternatively, distinct conformational dynamics of each paralog on each membrane may be essential for efficient membrane fusion. We now have the tools in-hand to test these models. Additionally, we can define changes in conformational dynamics associated with mitochondrial fusion defects and/or disease. For example, CMT2A variants with dysfunctional lipid mixing have been previously characterized, including Mfn1-F202L and Mfn1-S329P (Engelhart and Hoppins, 2019; Sloat and Hoppins, 2023). In the context of mini-Mfn1, these variants do not have altered GTPase activity, and both can dimerize via the GTPase domain. Full-length Mfn1-F202L is capable of fusion, albeit at a decreased efficiency to the native Mfn1. In contrast, while Mfn1-S329P initiates fusion, it is incapable of facilitating lipid mixing. Identifying the conformational dynamics of these variants will provide key information to connect conformational dynamics with steps in the membrane fusion process.

This would combine the protein structure, catalytic cycle, and the fusion mechanism into one concrete model. Finally, applying the system to full-length Mfn1 and Mfn2 would provide a more complete picture of allosteric regulation in the context of every domain and the membrane.

## Methods

### Constructs and Mutagenesis

The mini-Mfn1 construct was made by inserting the Mfn1-IM sequence from the Mfn1_IM_C pET28 plasmid (Yan et al., 2018; Engelhart and Hoppins, 2019) into the pETM11 vector (Uniprot accession #E0R11) followed by a GSSG linker, two TEV protease cleavage sites and a Twin-Strep-tag sequence using Gibson assembly. All mutations were made by Gibson assembly. Specifically, Mfn1-NoC changed six native cystines to create Mfn1-C111S-C156A-C167A-C196A-C260S-C327A-IM. The amber stop codon (TAG) replaced L13 to allow for integration of Acd. To generate a construct with both a donor and acceptor site, a single cysteine mutation was introduced into the donor-only construct with no native cystines at site Mfn1_NoC_-L13TAG-R171C-IM.

The following plasmids were used in this study for mammalian cell culture: pBABE-puro (#1764; Addgene), pBABE-mNG-Mfn1(M.m)-puro (Sloat and Hoppins, 2023). To create pBABE-mNG-Mfn1(H.s), the *Homo sapiens MFN1* sequence from Addgene #195162 was inserted into our pBABE vector with mNG at the N-terminus using Gibson assembly. pBABE-mNG-Mfn1(H.s)-NoC was created by replacing the DNA sequence for the first 364 residues of Mfn1 (H.s) with the sequence from pETM11-Mfn1-C111S-C156A-C167A-C196A-C260S-C327A-IM-GSSG-TEV-TwinStrep using Gibson assembly. After digestion with DpnI to remove template DNA, the amplified DNA was transformed into Endura (pBABE) or DH5-α (pETM11) *Escherichia coli* cells and plasmids were purified from selected colonies. All plasmids were confirmed by sequence analysis.

### Expression and Purification of the Mfn1-IM construct

All Mfn1-IM constructs described were co-transformed with the AcdA9 aminoacyl tRNA synthetase/tRNA-containing plasmid (pDule2)(Sungwienwong et al., 2017) into BL21 *E. coli* cells. Cells were then cultured in Luria-Bertani medium at 37°C with 50 μg/ml kanamycin and 60 μg/ml spectinomycin to an OD_600_ of ∼0.8. Protein expression was induced by adding β-D-1-thiogalactopyranoside (IPTG)(final concentration of 0.1 mM) and Acd (for a final concentration of 0.3 mM). These cultures were grown overnight (17-19 hrs) at 17.5°C. Cells were harvested by centrifugation, frozen in liquid nitrogen, and stored at −80°C. Cells were thawed with a room temperature water bath and resuspended in lysis buffer (30 mM Tris pH 7.4, 400 mM NaCl, 50 mM MgCl_2_, 2.5 mM β-mercaptoethanol) supplemented with HALT protease inhibitor cocktail.

Cell suspensions were lysed using Sonic Dismembrator Model 100 sonicator at power 5 for 4 seconds. This was repeated 3 times with the samples resting on ice for a minute between each cycle. The lysate was subject to centrifugation at 21,000 x *g* for 20 minutes. The supernatant was loaded onto 0.25 mL bed volume Strep-Tactin Superflow high-capacity beads (iba Biosciences) in a disposable gravity-flow column. The resin bound to protein was washed with 5 column volumes of wash solution (30 mM Tris pH 7.4, 400 mM NaCl, 50 mM MgCl_2_, 1 mM β-mercaptoethanol) and eluted with 625 μM d-Desthiobiotin (Sigma) in wash buffer. Mfn1-IM-containing elutions were treated with 100 μM tris(2-carboxyethyl) phosphine (TCEP) to fully reduce cysteines. Pure protein was desalted into KMgT buffer (50 mM KCl, 5 mM MgCl_2_, 30 mM Tris-HCl, pH 7.4, 500 mM TCEP) with a Zeba Spin Desalting Column (0.5 mL). The protein concentration was adjusted to 14 μM and then glycerol was added to 20%, for a final protein concentration of 10 μM, before the protein was aliquoted flash frozen with liquid nitrogen and stored at −80°C.

### Nucleotides and Labeling Reagents

Guanosine triphosphate (GTP) and guanosine 5’-diphosphate (sodium salt hydrate, GDP) were purchased from Sigma-Aldrich. Guanosine 5’-O-[gamma-thio] triphosphate (GTPγS) and guanosine 5’-[β,γ-imido] triphosphate (GMP-PNP) were both purchase from Cayman Chemical. GTP, GDP, and GTPγS were resuspended in 1M PIPES-KOH pH 7.6 (75 mM stocks). GMP-PNP was resuspended in 1M PIPES-KOH pH 7.6 (37.5 mM stock). [Cu(cyclenM)]^2+^ for tmFRET was prepared as previously described (Zagotta et al., 2021; Gordon et al., 2016).

### GTPase assay

Flash-frozen protein stored at −80°C was thawed on ice and diluted to a final concentration of 4.5 μM in reaction buffer (30 mM Tris-HCl pH 7.4, 50 mM KCl, 5 mM MgCl_2_, 100 μM TCEP). A 96-well plate was placed on ice and reactions were set up in triplicate. The indicated [GTP] was added to bring the reaction volume to 50 μL and reactions were incubated at 37°C for 15 min. To stop the reaction, the 96 well plate was moved to ice EDTA was added to a final concentration of 200 mM. Concentrations of free organic phosphate (Pi) were quantified with the malachite green reagent, which was incubated with the reaction for 15 minutes at room temperature. The absorbance at 650 nm was measured using a SpectraMax Plus 384 plate reader. A standard curve was generated with potassium phosphate and was used to determine the concentration of Pi resulting from GTP hydrolysis in each reaction. The data were analyzed with a Welch’s unpaired t-test.

### Cell culture and transfection

All cells were grown at 37°C and 5% CO_2_ and cultured in DMEM (Thermo Fisher Scientific) containing 1× GlutaMAX (Thermo Fisher Scientific) with 10% FBS (Seradigm). Mfn1-null MEF cells were purchased from the American Type Culture Collection.

### Retroviral transduction

Plat-E cells (Cell Biolabs) were maintained in complete media supplemented with 1 μg/ml puromycin and 10 μg/ml blasticidin and plated at ∼80% confluency the day before transfection. 350,000 Plat-E cells were plated in a six-well dish and the following day were transfected with pBABE plasmids (3 μg pBABE Mfn1 or an empty vector) using FuGENE HD (Promega) and Opti-MEM reduced serum media (Gibco). Transfection reagent was incubated overnight before a media change. Viral supernatants were collected at ∼48 and 72 hr post-transfection, passed through a 0.45 micron PES membrane, and incubated with MEFs in the presence of 8 μg/ml polybrene. Approximately 24 h after the last viral transduction, MEF cells were split, and selection was added (1 μg/ml puromycin).

### Microscopy

All cells were plated on No. 1.5 glass-bottomed dishes (MatTek) approximately 48 hr before imaging. Cells were incubated with 0.1 μg/ml MitoTracker Red CMXRos for 15 min at 37°C with 5% CO_2_. MEF cells were then rinsed into complete media and incubated for at least one hour before addition of one drop of NucBlue (Molecular Probes). A Z-series with a step size of 0.25 μm was collected with a Nikon Ti2E confocal microscope with a 63X NA 1.4 (numerical aperture) oil objective (Nikon), a Cicero spinning disk, using a 5 channel Celesta light source (Nikon), and an sCMOS camera (Prime BSI Express). Each cell line was imaged on at least three separate occasions by a blinded experimenter (n > 100 cells per experiment).

### Image analysis

Images were deconvolved using 15 iterations of 3D Landweber deconvolution. Deconvolved images were analyzed using Nikon Elements software. Maximum intensity projections were created using ImageJ software (NIH). Mitochondrial morphology in mammalian cells was scored as follows: hyperfused indicates that the entire mitochondrial network in the cell was connected as a single structure; reticular indicates that fewer than 30% of the mitochondria in the cell were fragments (fragments defined as mitochondria less than 2.5 μm in length); fragmented indicates that most of the mitochondria in the cell were less than 2.5 μm in length. In MEF cells expressing mNeonGreen-Mfn1, only cells with GFP signal were scored.

### TCSPC Lifetime measurements

TCSPC lifetime data of Acd labeled protein samples at 300 nM (100 μL) was measured in 50 μL volume quartz cuvettes (Starna Cells, inc). Lifetime data were measured using a PicoQuant FluoTime 300 Fluorescence Lifetime Spectrometer (PicoQuant, Berlin Germany) with 375 nm UV laser excitation and 446 nm emission. Single photo arrivals were recorded on a hybrid photomultiplier detector assembly (PMA-40). Four cuvettes were consecutively recorded, the first with Mfn1-L13Acd-R171C, the second with Mfn1-L13Acd-NoC, the third with buffer-only correspondent to the buffer of each experiment, and the fourth with diluted LUDOX in MiliQ water for measurement of the instrument response function (IRF). First the lifetime of all cuvettes was recorded without the labelling of [Cu(cyclenM)]^2+^. Then 1 μL of 11 mM [Cu(cyclenM)]^2+^ was added and the lifetimes were measured until labelling was stable. This was done in the following buffer conditions:

Apo: 50 mM KCl, 5 mM MgCl_2_, 30 mM Tris-HCl, pH 7.4
GDP: 50 mM KCl, 5 mM MgCl_2_, 30 mM Tris-HCl, pH 7.4, 750 μM GDP
GMP-PNP: 50 mM KCl, 5 mM MgCl_2_, 30 mM Tris-HCl, pH 7.4, 750 μM GMP-PNP
GTPγS: 50 mM KCl, 5 mM MgCl_2_, 30 mM Tris-HCl, pH 7.4, 750 μM GTPγS
GDP•BeF_3_: 50 mM KCl, 5 mM MgCl_2_, 30 mM Tris-HCl, pH 7.4, 750 μM GDP, 25 mM NaF, 250 mM BeSO_4_

### FRET model of time-domain fluorescence lifetime data to obtain Gaussian distance distributions

Fitting of the fluorescence lifetime data was as previously described (Eggan et al., 2025). Briefly:

The time-domain fluorescence lifetime data of the donor fluorophore in the presence of the acceptor were fit by model estimates for the time course of the decay, *Decay*(*t*). These fits were calculated from the convolution of the measured instrument response function, *IRF*(*t*) of the system with model estimates of the fluorescence lifetime of the donor in the presence of the acceptor, *I*_*DA*_(*t*) and buffer-only fluorescence *I*_*B*_(*t*):

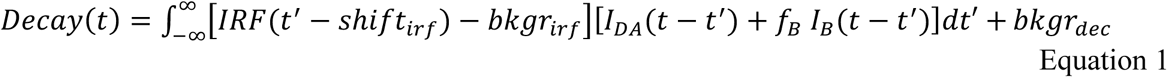

Where the variables are defined in Table I and displayed graphically in Fig. S3.

**Table I.**

| Variable | Description |
| --- | --- |
| $t$ | Time of photon emission (in s) |
| $r$ | Distance between the donor and acceptor (in Å) |
| $Decay(t)$ | Time-dependent decay of the donor fluorescence (in counts) |
| $DecayB(t)$ | Time-dependent decay of the buffer-only fluorescence (in counts) |
| $I_{DA}(t)$ | Model estimate of the fluorescence lifetime of the donor in the presence of the acceptor (in counts) |
| $I_B(t)$ | Model estimate of the fluorescence lifetime of a buffer-only sample (in counts) |
| $IRF(t)$ | Measured instrument response function (in counts) |
| $bkgr_{dec}$ | Time-independent decay background (in counts) |
| $bkgr_{irf}$ | Time-independent background of the instrument response function (in counts) |
| $shift_{irf}$ | Time shift between the instrument response function and the measured decay (in ps) |
| $f_B$ | Scaling of the buffer fluorescence |
| $A_0$ | Amplitude of the model estimate of the fluorescence lifetime of the donor (in counts) |
| $f_D$ | Fraction of donor-only fluorescence in the sample |
| $\alpha_{Di}$ | Fraction of the $i$th component of the donor-only fluorescence (in counts) |
| $\tau_{Di}$ | Time constant of the $i$th component of the donor-only fluorescence (in s) |
| $\alpha_{Bi}$ | Amplitude of the $i$th component of the buffer fluorescence (in counts) |
| $\tau_{Bi}$ | Time constant of the $i$th component of the buffer fluorescence (in s) |
| $R_0$ | Characteristic distance between donor and acceptor producing 50% FRET efficiency |
| $\rho(r)$ | Probability distance distribution of the donor and acceptor distances |
| $f_{Ai}$ | Fraction of the $i$th component of the probability distance distribution |
| $\bar{r}_i$ | Average distance of the $i$th component of the donor-only fluorescence (in Å) |
| $\sigma_i$ | Standard deviation of the $i$th component of the donor-only fluorescence (in Å) |

The model for fluorescence lifetime assumes a donor-only fluorescence lifetime with one or two exponential components, and FRET between a donor and acceptor separated by one or two Gaussian-distributed distances. This model predicts the following relationship for the fluorescence lifetime of the donor in the presence of acceptor, *I*_*DA*_(*t*):

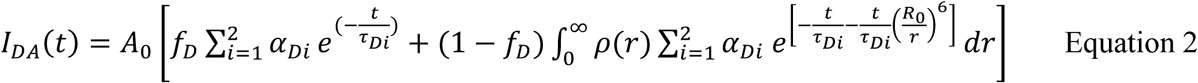

Where *⍺*_*D*1_ + *⍺*_*D*2_ = 1.

The density distribution of donor-acceptor distances, *ρ*(*r*), was assumed to be the sum of up to two Gaussians:

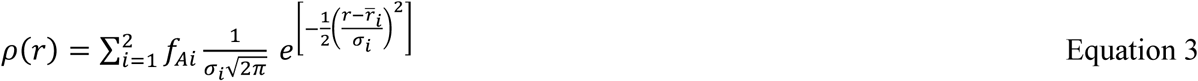

where *f*_*A*1_ + *f*_*A*2_ = 1.

The decay time course of the buffer-only sample, *DecayB*(*t*), was fit with a convolution of the measured instrument response function, *IRF*(*t*) with a multi-exponential model for the fluorescence lifetime, *I*_*B*_(*t*):

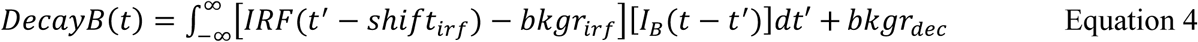

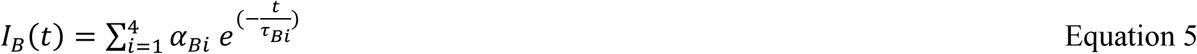

This model for the buffer-only fluorescence lifetime was then added to the model for the sample lifetime before convolving with the *IRF*(*t*) (Equation 1).

This FRET model for fluorescence lifetimes and FRET was implemented in Igor Pro v8 (Wavemetrics, Lake Oswego, OR) (code available at https://github.com/zagotta/TDlifetime_program). The FRET model was fit, with *χ*^2^ minimization, assuming a standard deviation equal to 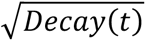, to individual decay time courses. There are 15 parameters in the parameter vector (*f*_*D*_, *τ*_*D*1_, *⍺*_*D*1_, *τ*_*D*2_, *R*_0_, *r̅*_1_, *σ*_1_, *f*_*A*2_, *r̅*_2_, *σ*_2_, *shift*_*irf*_, *f*_*B*_, *A*_0_, *bkgr*_*dec*_, *bkgr*_*irf*_). *R*_0_ was fixed to a value previously determined from emission spectra of the donor and absorbance spectra of the acceptor using the Förster equation assuming ***κ***^2^ = 2/3. *f*_*A*2_,was generally fixed to 0 or 1 for fits assuming a single Gaussian distance distribution but was allowed to vary when multiple distance components were considered (e.g. in the GDP•BeF_3_ condition). *f*_*B*_ was fixed to the acquisition time for the experimental sample divided by the acquisition time for the buffer-only sample. And *bkgr*_*irf*_was always fixed to 0 as the average background photon count in the *IRF*(*t*) was <<1.

For the individual decay fits to the donor-only sample, *f*_*D*_ was set to 1, and only *τ*_*D*1_, *⍺*_*D*1_, *τ*_*D*2_, *shift*_*irf*_, *A*_0_, *bkgr*_*dec*_ were allowed to vary. For the individual decay fits to the donor+acceptor samples, *τ*_*D*1_, *⍺*_*D*1_, *τ*_*D*2_were constrained to the previously determined values from the donor-only sample, and *f*_*D*_, *r̅*_1_, *σ*_1_, *f*_*A*2_, *r̅*_2_, *σ*_2_, *shift*_*irf*_, *A*_0_, *bkgr*_*dec*_ were varied.

## Supporting information

https://submit.biorxiv.org/submission/submit?msid=BIORXIV/2025/681278&gotoPage=1&nextpage=guide&guidehelp=true

## Acknowledgements

We thank the Oregon State University GCE4ALL (Center for Genetic Code Expansion for All) for their long-standing collaboration, Dr. James Peterson and Dr. Kyle Shaffer for sharing Acd, and Dr. Pierce Eggan, Charlie DeFreest, Dr. Shauna Otto, and Dr. Stephanie Sloat for technical support and scientific discussions. Funding was provided by grants from the National Institute of General Medical Sciences (R01GM118509 to SH; TGGM153507 to SMH; R35GM145225 to SEG; R35GM148137 to WNZ). The authors declare no competing financial interests.

## Author contributions

Sophie Hurwitz: conceptualization, data curation, formal analysis, funding acquisition, investigation, methodology, validation, visualization, and writing—original draft, review, and editing. William Zagotta: formal analysis, funding acquisition, investigation, methodology, resources, validation, visualization, and writing—editing. Sharona Gordon: formal analysis, funding acquisition, investigation, methodology, resources, validation, visualization, and writing—editing. Suzanne Hoppins: conceptualization, data curation, formal analysis, funding acquisition, investigation, methodology, project administration, resources, supervision, validation, visualization, and writing—original draft, review, and editing.

## Supplemental Figure Legends

**Supplemental Figure 1:**
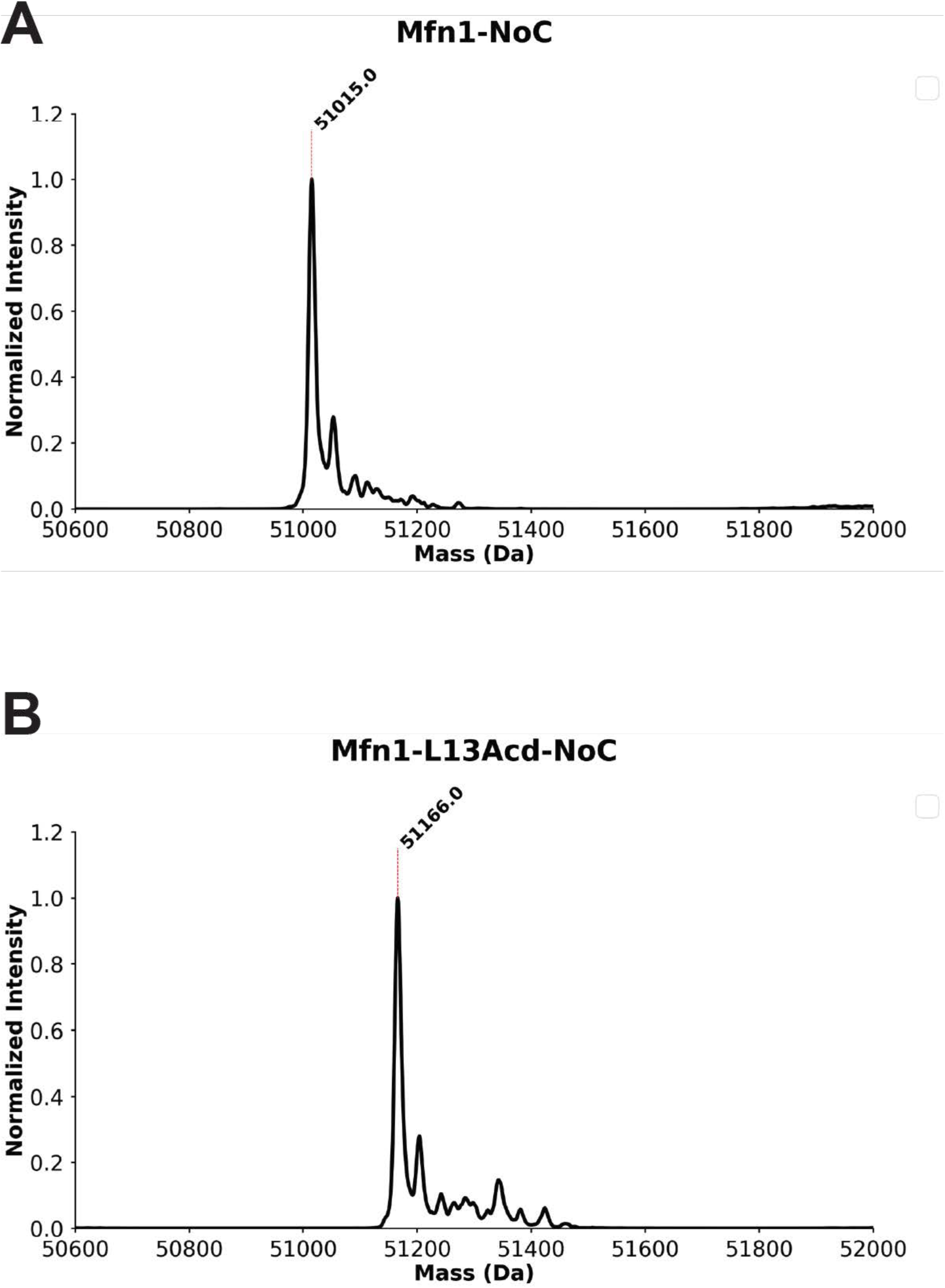
Acd Incorporation at L13 of mini-Mfn1. **(A)** Mass spectrometry analysis of mini-Mfn1-NoC to get the predominate product off the column of 51015.0 Da, which agrees with the mass of the mini-Mfn1-NoC construct without the initial methionine, which is commonly cleaved in the cytoplasm of BL21 *E. coli* cells (Wingfield, 2017). There are no other comparable peaks, demonstrating a clean purification**. (B)** Mass Spectrometry analysis of mini-Mfn1-L13Acd-NoC to get the predominate product of 51166.0 Da. This demonstrates a change in 151 Da from Acd incorporation, which only arises from a leucine being substituted for Acd. The absence of other comparable peaks demonstrates specific integration.

**Supplemental Figure 2:**
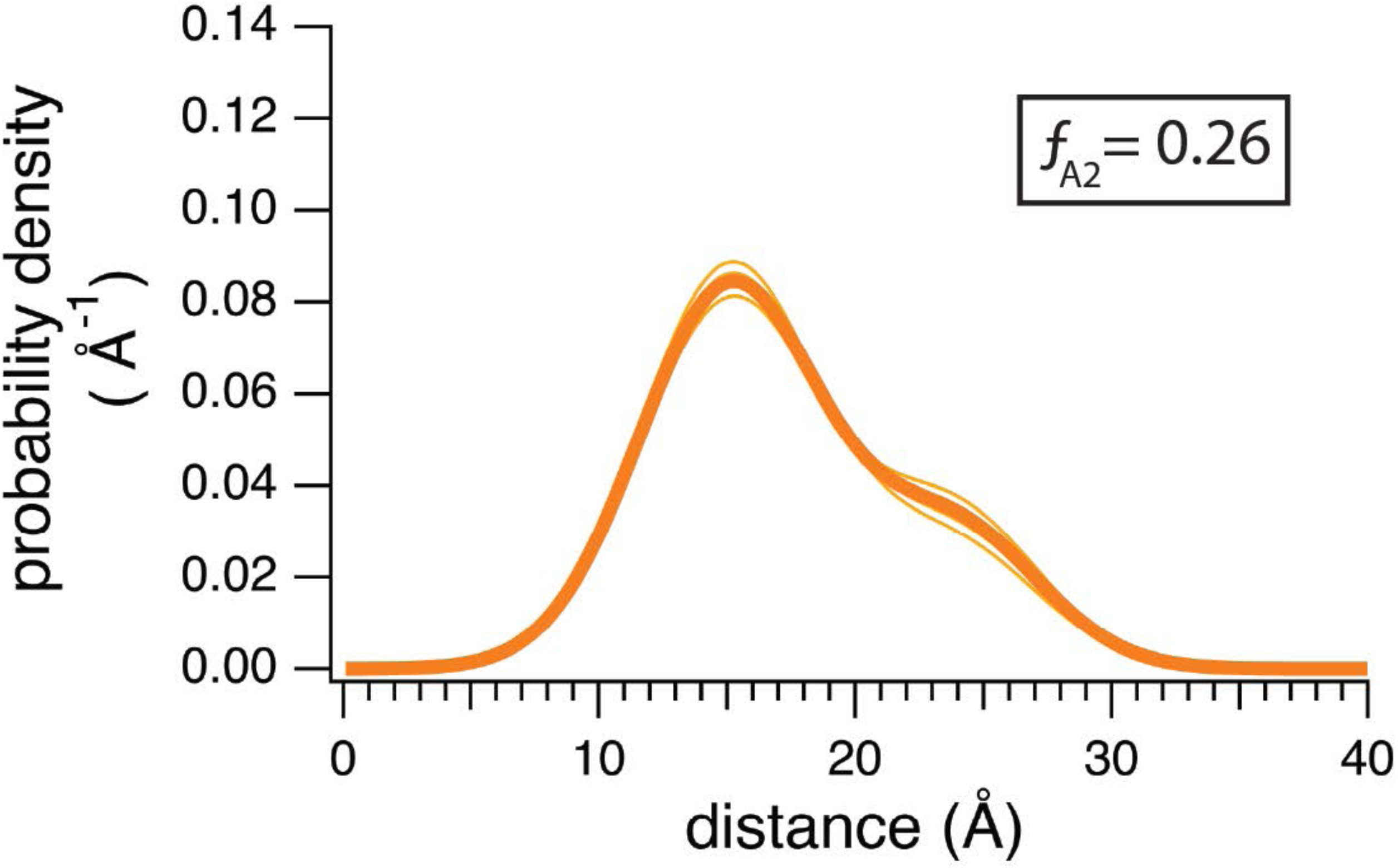
Steady-state lifetime tmFRET of mini-Mfn1 in the apo state fit to a two-state model. Spaghetti plot of distributions from individual experiments in the absence of nucleotide fit to a dual Gaussian with the average in bold. The fraction in the closed state (ƒ_A2_) is 0.26. The reduced chi-square values for each of the individual replicates are as follows: rep1=0.68, rep 2=0.62, rep 3=0.74, rep 4=0.71, rep 5=0.70, rep 6=0.70.

**Supplemental Figure 3:**
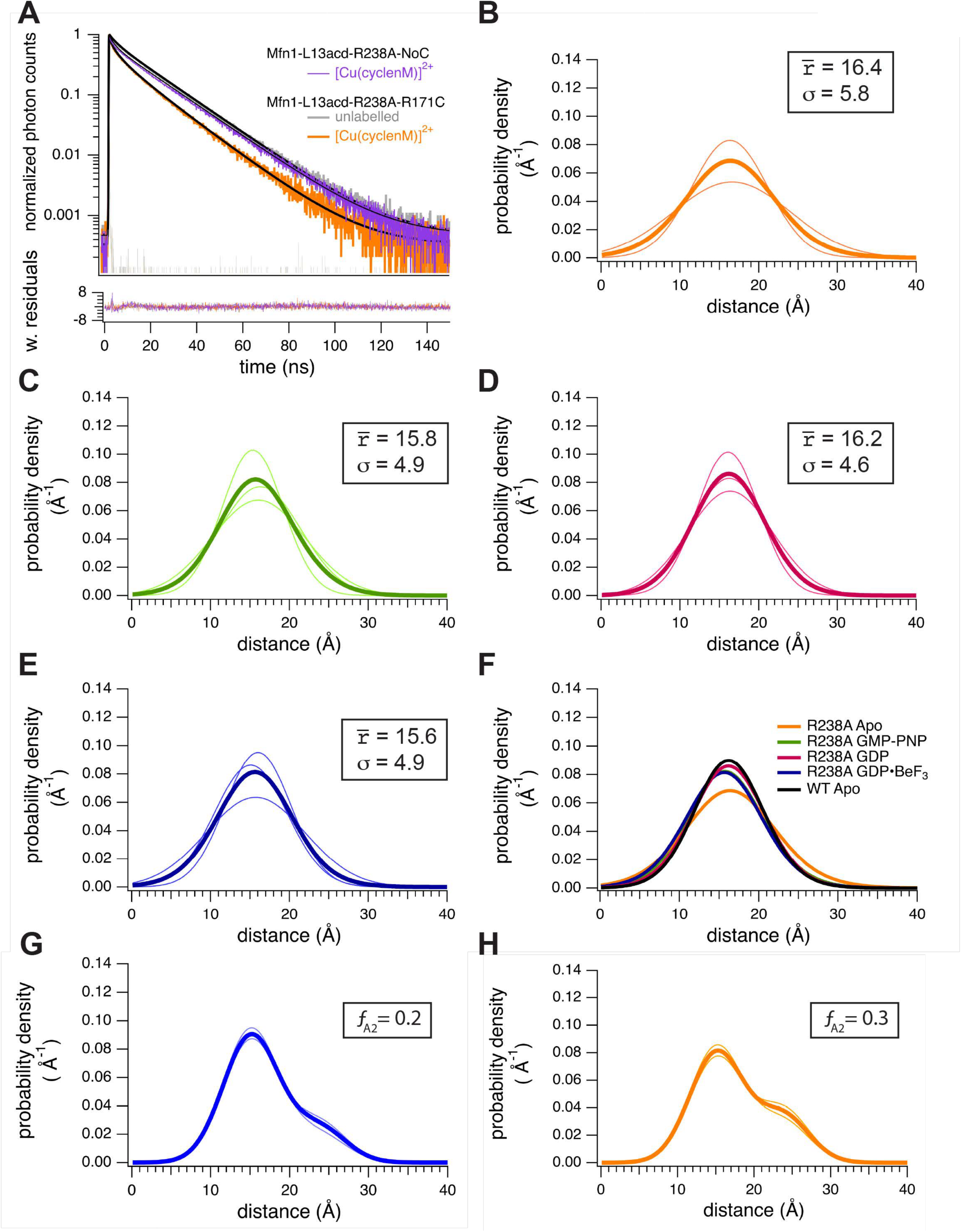
Mfn1-L13Acd-R171C-R238A resembles the apo state in all stages of the catalytic cycle. (**A**) Representative TCSPC data from Mfn1-L13Acd-R238A-NoC with no nucleotide in the presence of [Cu(cyclenM)]^2+^(violet) and Mfn1-L13Acd-R238A R171C before (grey) and after (orange) the addition of [Cu(cyclenM)]^2+^. (**B**) Spaghetti plot of distributions from individual experiments with no nucleotide bound with the average of the waves displayed in bold. (**C**) Spaghetti plot of distributions from individual experiments with GMP-PNP bound with the average of the waves displayed in bold. (**D**) Spaghetti plot of distributions from individual experiments with GDP bound with the average of the waves displayed in bold. (**E**) Spaghetti plot of distributions from individual experiments with GDP•BeF_3_ bound with the average of the waves displayed in bold. (**E**) Plot of average wave distributions for apo (orange), GMP-PNP (green), GDP (magenta) GDP•BeF_3_ (blue), and apo-WT-Mfn1 (black). (**F**) Spaghetti plot of distributions from individual experiments in the absence of nucleotide fit to a dual Gaussian with the average in bold. The fraction in the closed state (ƒ_A2_) is 0.20. The reduced chi-square values for each of the individual replicates are as follows: rep1=1.01, rep 2=0.65, rep 3=0.79. (G) Spaghetti plot of distributions from individual experiments in GDP•BeF_3_ fit to a dual Gaussian with the average in bold. The fraction in the closed state (ƒ_A2_) is 0.3. The reduced chi-square values for each of the individual replicates are as follows: rep1=0.88, rep 2=0.79, rep 3=0.94.

**Supplemental Figure 4.**
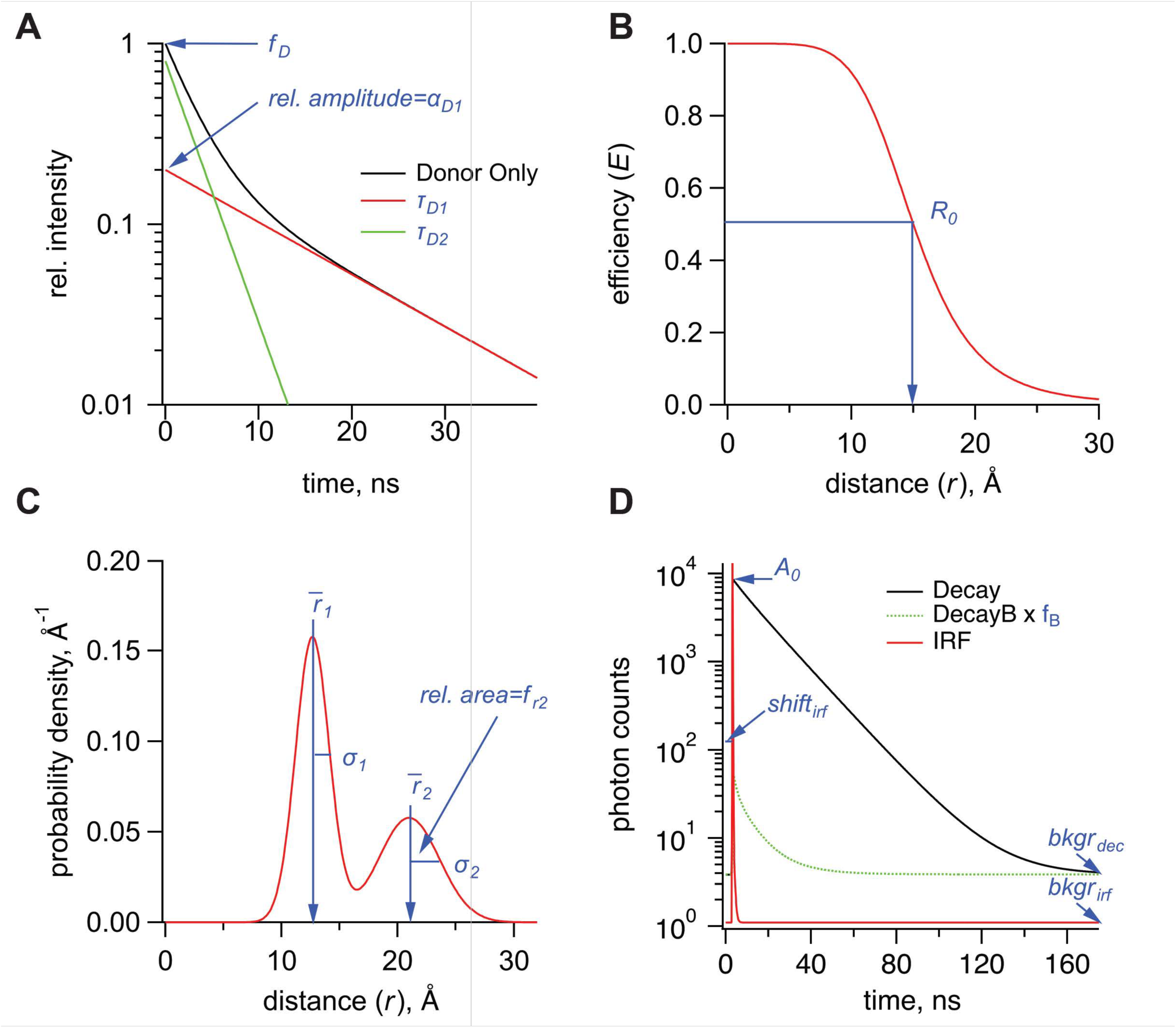
Parameters used in the FRET model for TCSPC data. **(A)** Graph of donor-only fluorescence-lifetime with two exponential components with time constants (*τ*_*D*1_, *τ*_*D*2_) with relative amplitude *⍺*_*D*1_. The amplitude fraction of the donor-only lifetime in an experiment is determined as *f*_*D*_. (**B)** FRET efficiency (*E*) plot as a function of distance (*r*) between donor and acceptor and the *R_0_* values for 50% FRET transfer. **(C)** Probability distribution plot of donor and acceptor distances P(*r*) showing the sum of two Gaussian distributions, each with their own average distance (*r*R_1_ and *r*R_2_), standard deviations (*σ*_1_ and *σ*_2_) and relative amplitude of the second component (*f*_*r*2_). **(D)** Example TCSPC data is shown with experimental measured photon count data (Decay), the instrument response function (IRF), and the buffer only decay (DecayB), with its corresponding scaling factor *f*_*B*_ relative to the Decay trace. Also indicated is a time-independent background of photon counts for the Decay trace, *bkgr*_*dec*_, as well as two parameters associated with the IRF, its background, *bkgr*_*irf*_, and the shift between instrument response function and measured decay, *shift*_*irf*_. Parameter *A*_0_ is the amplitude of the FRET model estimate of the fluorescence lifetime of the donor (in photon counts).

