## Supplementary material for "Time-resolved tmFRET reveals GTP-coupled conformational changes in Mfn1": https://submit.biorxiv.org/submission/submit?msid=BIORXIV/2025/681278&gotoPage=1&nextpage=guide&guidehelp=true

Author Affiliations:
a Department of Biochemistry, University of Washington, Seattle, WA 98195
b Department of Neurobiology and Biophysics, University of Washington, Seattle, WA 98195

Corresponding author:
Suzanne Hoppins
University of Washington
Department of Biochemistry
1959 NE Pacific Street
Health Sciences Building, J383
Seattle, WA 98195

ORCID – orcid/org/0000-0002-8070-3560

Keywords: allostery, dynamin superfamily protein, tmFRET, GTPase

Fig S1

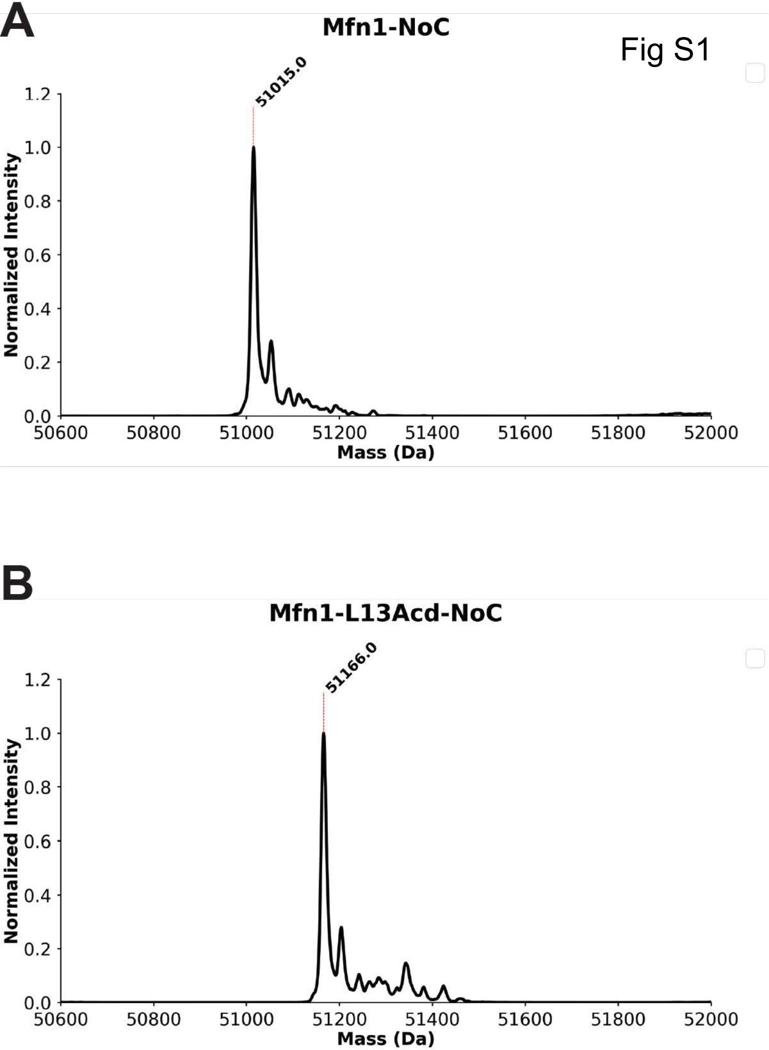

A
Mfn1-NoC
51015.0
Normalized Intensity
1.2
1.0
0.8
0.6
0.4
0.2
0.0
50600
50800
51000
51200
51400
51600
51800
52000
Mass (Da)
B
Mfn1-L13Acd-NoC
51166.0
Normalized Intensity
1.2
1.0
0.8
0.6
0.4
0.2
0.0
50600
50800
51000
51200
51400
51600
51800
52000
Mass (Da)

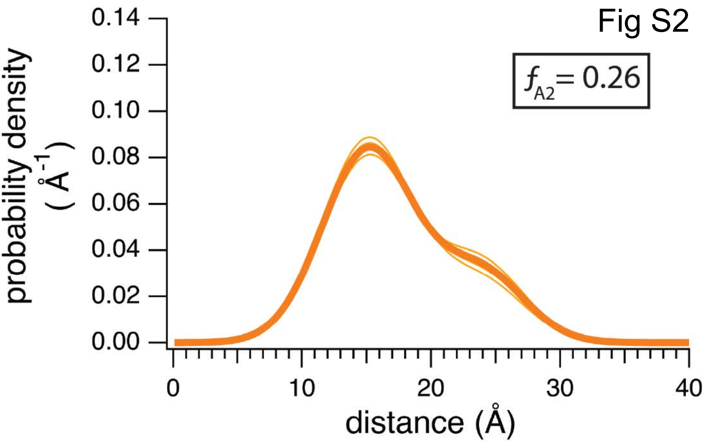

Fig S2
$f_{A2}$= 0.26
probability density (Å$^{-1}$)
0.14
0.12
0.10
0.08
0.06
0.04
0.02
0.00
0
10
20
30
40
distance (Å)

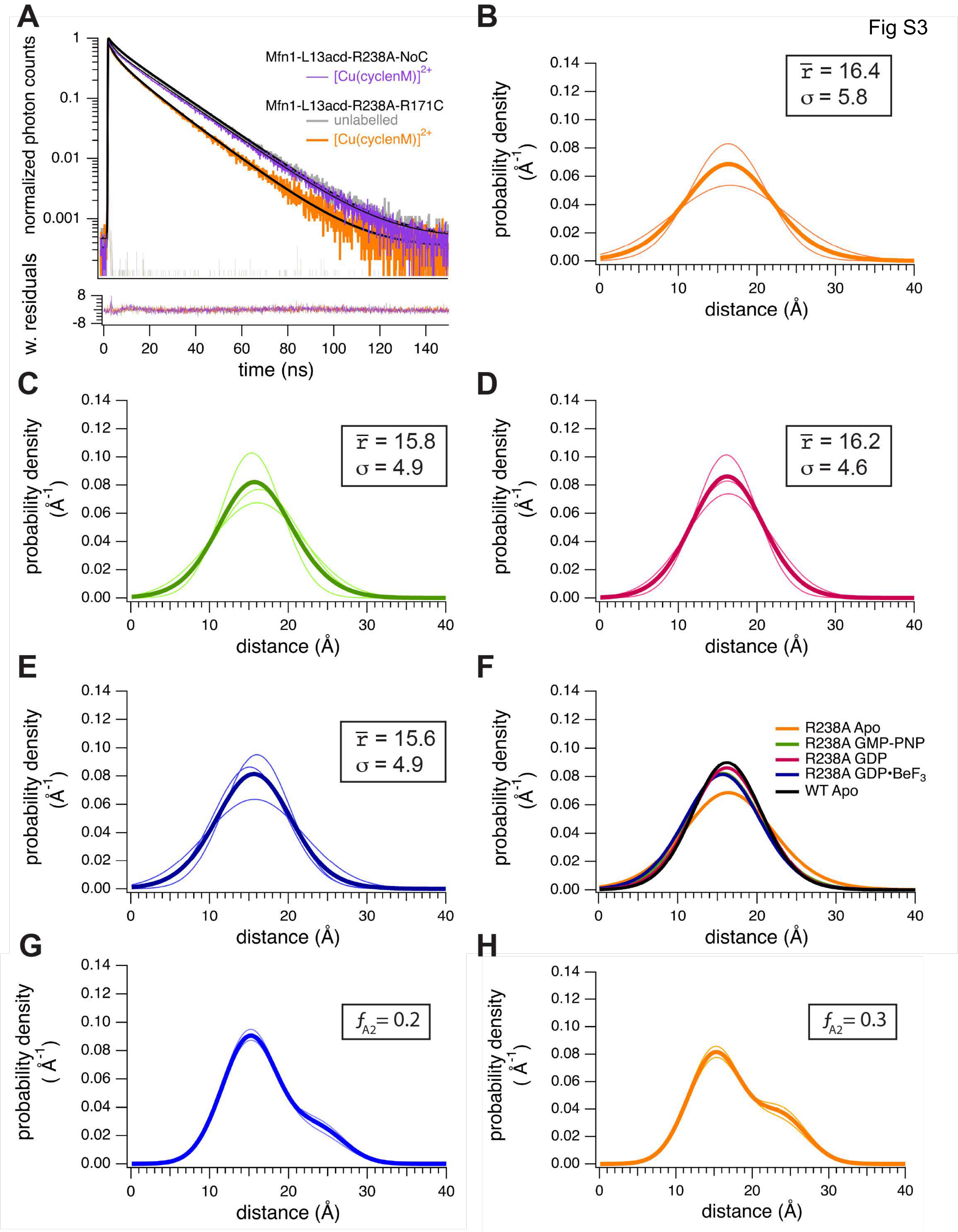

A
B
C
D
E
F
G
H
Fig S3
normalized photon counts
w. residuals
time (ns)
Mfn1-L13acd-R238A-NoC
[Cu(cyclenM)]2+
Mfn1-L13acd-R238A-R171C
unlabelled
[Cu(cyclenM)]2+
probability density (Å-1)
distance (Å)
r̄ = 16.4
σ = 5.8
r̄ = 15.8
σ = 4.9
r̄ = 16.2
σ = 4.6
r̄ = 15.6
σ = 4.9
R238A Apo
R238A GMP-PNP
R238A GDP
R238A GDP•BeF3
WT Apo
fA2 = 0.2
fA2 = 0.3

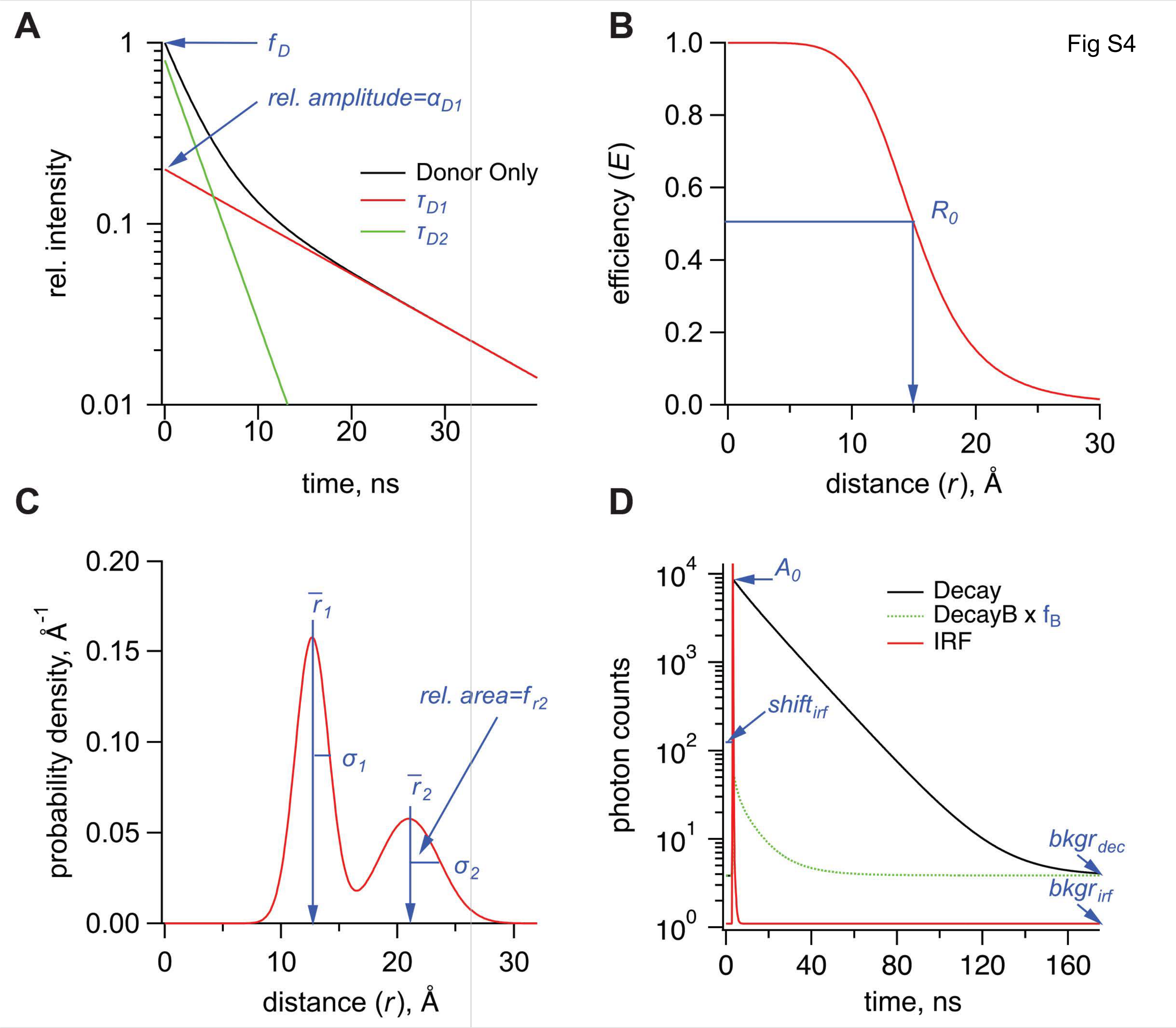

A
B
C
D
Fig S4
1
0.1
0.01
0
10
20
30
rel. intensity
time, ns
f_D
rel. amplitude=α_D1
Donor Only
τ_D1
τ_D2
1.0
0.8
0.6
0.4
0.2
0.0
efficiency (E)
distance (r), Å
R_0
0.20
0.15
0.10
0.05
0.00
probability density, Å^-1
r̄_1
σ_1
r̄_2
σ_2
rel. area=f_r2
10^4
10^3
10^2
10^1
10^0
photon counts
time, ns
0
40
80
120
160
A_0
Decay
DecayB x f_B
IRF
shift_irf
bkgr_dec
bkgr_irf
